# A flux-based machine learning model to simulate the impact of pathogen metabolic heterogeneity on drug interactions

**DOI:** 10.1101/2021.08.03.454957

**Authors:** Carolina H. Chung, Sriram Chandrasekaran

**Author notes:** **Author contributions** Conceptualization: SC Methodology: CHC, SC Investigation: CHC, SC Visualization: CHC Supervision: SC Writing—original draft: CHC Writing—review & editing: CHC, SC.

## Abstract

Drug combinations are a promising strategy to counter antibiotic resistance. However, current experimental and computational approaches do not account for the entire complexity involved in combination therapy design, such as the effect of pathogen metabolic heterogeneity, changes in the growth environment, drug treatment order and time interval. To address these limitations, we present a comprehensive approach that uses genome-scale metabolic modeling and machine learning to guide combination therapy design. Our mechanistic approach (a) accommodates diverse data types, (b) accounts for time- and order-specific interactions, and (c) accurately predicts drug interactions in various growth conditions and their robustness to pathogen metabolic heterogeneity. Our approach achieved high accuracy (AUC = 0.83 for synergy, AUC = 0.98 for antagonism) in predicting drug interactions for *E. coli* cultured in 57 metabolic conditions based on experimental validation. The entropy in bacterial metabolic response was predictive of combination therapy outcomes across time scales and growth conditions. Simulation of metabolic heterogeneity using population FBA identified two sub-populations of *E. coli* cells defined by the levels of three proteins (eno, fadB and fabD) in glycolysis and lipid metabolism that influence cell tolerance to a broad range of antibiotic combinations. Analysis of the vast landscape of condition-specific drug interactions revealed a set of 24 robustly synergistic drug combinations with potential for clinical use.

**Significance:** Worldwide, 700,000 people die each year from drug-resistant infections. Drug combinations have great potential to reduce the spread of drug-resistant bacteria. However, their potency is impacted by both the pathogen growth environment and the heterogeneity in pathogen metabolism. The metabolic heterogeneity in a pathogen population allows them to survive antibiotic treatment. Here we present a flexible machine-learning framework that utilizes diverse data types to effectively search through the large design space of both sequential and simultaneous combination therapies across hundreds of simulated growth conditions and pathogen metabolic states. Our approach can serve as a useful guide for the selection of robustly synergistic drug combinations.

## Introduction

Antimicrobial resistance (AMR) occurs due to extended exposure to antibiotics, which allows bacteria to evolve resistance mechanisms that render antibiotic treatments ineffective^1^. In the context of AMR, bacterial metabolism plays a key role. Cell-to-cell variation in metabolism within a population can be beneficial in responding to antibiotic stress^2,3^, and several pathogens take on a distinct metabolic state *in vivo* to tolerate antibiotics^4,5^. It is important to note that tolerant cells are predicted to be the source of drug-resistant pathogens^5–7^. In addition to stochasticity in metabolic activity within a population, extrinsic factors such as the metabolic environment also influence antibiotic efficacy^8–10^. For example, the availability of oxygen and extracellular metabolites modulate potency of antibiotics^9^. Metabolism can thus promote pathogen survival through adaptable use of nutrients in the local environment ^11,12^. Bacterial metabolism also impacts susceptibility to antibiotics through the production of reactive oxygen species^13,14^ or changes in membrane permeability^9^. Of note, these metabolic responses are also tied to entropy (i.e., disorder) in the bacterial stress response, which has been shown to be a generalizable predictor for antibiotic sensitivity^15^. Altogether, these individual findings suggest that modeling bacterial metabolism in response to antibiotics may be insightful for the design of novel treatments that mitigate resistance.

Combination therapy, which involves the use of two or more therapeutics, holds great potential for combating resistant pathogens as it not only leverages already regulated therapeutics^16^, but also offers room for improved efficacy^17^. Further, combination therapy could be optimized to selectively target resistant pathogens via collateral sensitivity, which has been shown to overcome multi-drug resistance in cancer^18,19^. Collateral sensitivity entails the increased sensitivity to a therapeutic that results from initial treatment with another stress agent^20^. This phenomenon has been observed across various diseases and organisms^18,21–23^, and in context of AMR, could be leveraged to prevent and mitigate resistance^24^. Theoretical studies have also predicted that antibiotic combinations can be effective in heterogeneous populations and reduce the rise of resistance more effectively than monotherapy^24,25^. However, these studies do not provide a guideline to identify promising combinations among thousands of possible candidates. Combination therapies are traditionally identified using experimental methods; however, this approach quickly becomes infeasible when considering the vast combinatorial search space, the effects of the growth media, pathogen metabolic heterogeneity, and time-/order-dependence for treatment efficacy.

With the advent of high-throughput omics data and application of machine learning (ML), it is now possible to expedite the search for effective combination therapies. ML has also been applied to reveal mechanistic insights into antibiotic mechanisms of action^26,27^ and identify novel antibacterial compounds^28,29^. In the past decade, several groups have used these methods to computationally design combination therapies in context of cancer^30–35^ and AMR^36–38^. For the latter case, prior models have been shown to generate predictions that accurately correspond to experimental and clinical efficacy against *Escherichia coli* and *Mycobacterium tuberculosis*, thus offering effective reduction of the search space for combination therapies against AMR^36,38^. However, these approaches are limited by the availability of omics data measuring the bacterial response to antibiotic treatment. The combined drug effect on bacterial growth has also only been assessed in a limited number of growth environments^37^. Moreover, current models have primarily focused on simultaneous combinations; consequently, the potential of designing time- and order-dependent combination therapies that promote collateral sensitivity remains unexplored. Since combination therapy is increasingly used to treat many medical conditions such as tuberculosis (TB), gram negative-, and biofilm associated-infections^39–42^, it is essential to consider how various metabolic factors (e.g., cell-to-cell heterogeneity, growth environment) influence the efficacy of different drug combinations. Computational tools are hence necessary to identify antibiotic treatments that are robustly efficacious across heterogeneous environments^43,44^.

To address these limitations, we present an approach that integrates genome-scale metabolic models (GEMs) into ML model development to determine effective combination therapies. Using GEMs allows us to integrate diverse data types and account for different pathogen metabolic states and growth conditions. GEMs are computational models built from gene-protein-reaction associations of metabolic genes present in the genome of an organism^45^. Additionally, they include annotation of traditional antibiotic targets such as cell wall synthesis, DNA replication, and RNA transcription. Model constraints, such as from omics data or nutrient availability, can be imposed to simulate bacterial metabolism in response to different perturbations^46,47^. Our approach using GEMs and ML provides a systems-level perspective of the bacterial response to antibiotic treatment in condition-specific cases. This is critical for designing efficacious combination therapies, since experimentally measured susceptibility to antibiotics may not always translate into efficacy *in vivo*. We further extend our approach to predict outcomes for sequential combination therapies, which can be designed into cyclic antibiotic regimens that mitigate resistance^24^. Finally, we showcase how our models reveal mechanistic insights that explain treatment potency and can be leveraged to finetune data-driven combination therapy design.

## Results

### The CARAMeL approach for combination therapy design

Our approach, called *Condition-specific Antibiotic Regimen Assessment using Mechanistic Learning* (*CARAMeL*), involves a two-step modeling process: (a) simulating metabolic flux data using GEMs and (b) developing a ML model to predict combination therapy outcomes using flux from GEMs. For the first part, omics data and metabolite composition of the extracellular environment serve as GEM inputs to determine flux profiles in response to drug treatment and growth in defined media, respectively (***Fig. 1***). For the second part, GEM-derived flux profiles and drug interaction data serve as inputs to train a ML model that predicts interaction outcomes for novel drug combinations (***Fig. 1***). We developed ML models predictive of combination therapy outcomes for *E. coli* and *M. tb* using the Random Forests algorithm. We specifically chose this ML method as it can handle small datasets and determine feature importance, i.e., how much each feature contributes to the accuracy in model predictions. The feature importance can reveal mechanistic insights into the factors driving combination therapy outcomes.

**Fig. 1.**
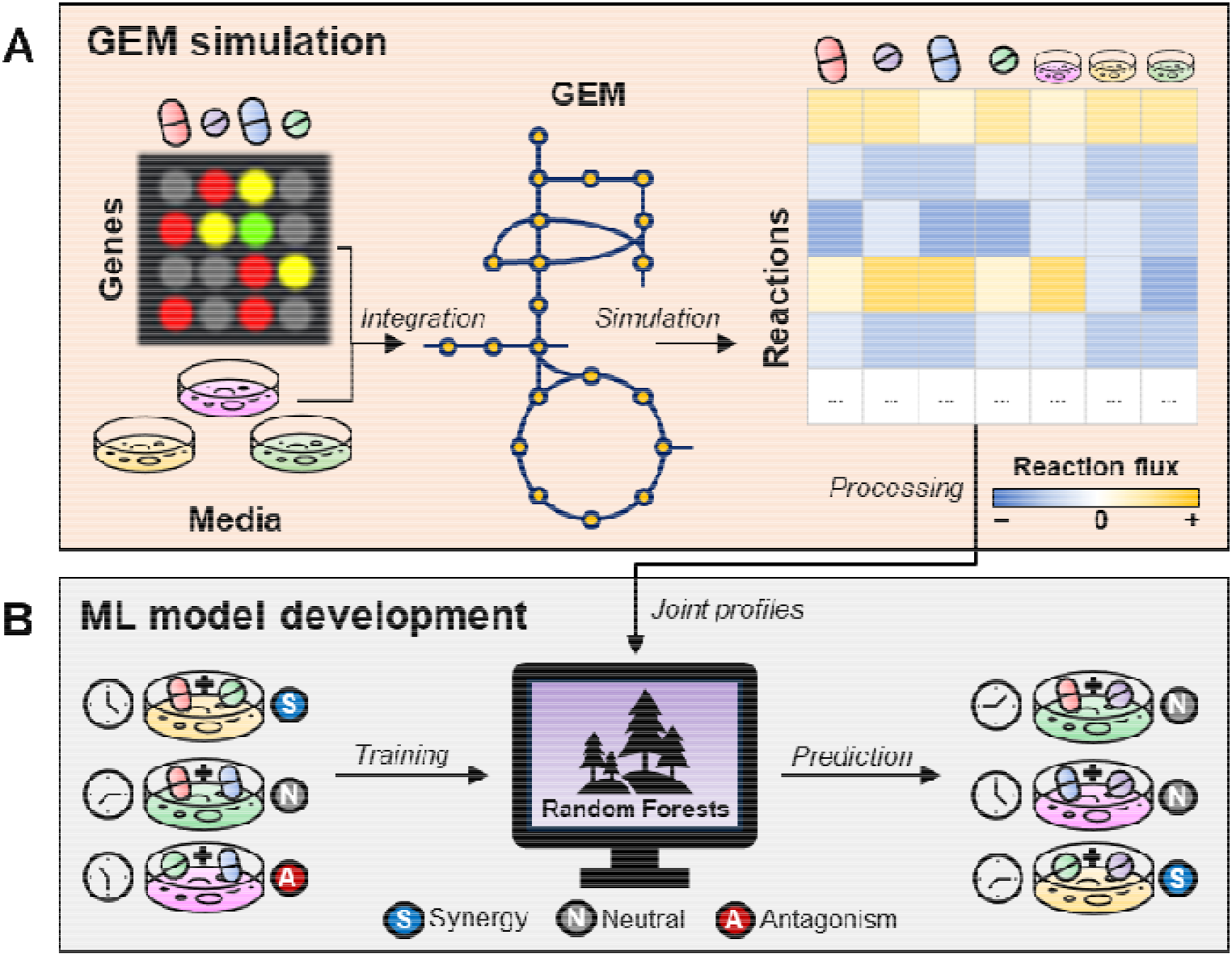
CARAMeL approach schematic. The *Condition-specific Antibiotic Regimen Assessment using Mechanistic Learning* (CARAMeL) approach involves a two-step process: (**A**) omics data (e.g., transcriptomics) measured for single drug treatments and information on growth media composition are integrated into a genome-scale metabolic model (GEM) to simulate metabolic flux changes. (**B**) This information, along with drug interaction data, serve as inputs to train a machine learning (ML) model; the trained model can then be used to predict outcomes for novel drug interactions.

We determined metabolic flux profiles in response to drug treatment and condition-specific growth by constraining the *E. coli* GEM iJO1366^48^ and the *M. tb* GEM iEK1011^49^. For drug flux profiles, we imposed chemogenomic data for *E. coli*^50^ and transcriptomic data for *M. tb*^38^ as GEM constraints. Briefly, chemogenomic data measures single-gene knockout (KO) fitness while transcriptomics data measures genome-wide expression of genes. By selecting genes for which there was differential fitness or expression in response to a specific treatment, we could infer a set of differentially regulated genes for individual drugs. For transcriptomic data, positive and negative differential expression directly corresponded with up- and down-regulation, respectively. For chemogenomic data, we assumed that gene KOs that result in low fitness are likely to be up-regulated upon drug treatment, while gene KOs that enhance fitness were likely to be down-regulated. This assumption is based on the cost-benefit gene expression model proposed by Dekel & Alon^51^. Direct comparison of flux profiles simulated from a chemogenomic-based approach against flux profiles simulated with transcriptomics and proteomics data confirmed that these assumptions were valid (***Fig. S1***)^52–54^. To determine growth media flux profiles, the availability of metabolites within a media condition was used to constrain the GEMs. Specifically, we modified the uptake rate for exchange reactions providing key metabolites (e.g., glycerol exchange for M9 glycerol media) to allow cellular intake (see ***Methods*** for further details).

Prior to ML model development, we processed drug and media flux profiles to determine joint profiles for all combinations of interest. Joint profiles are comprised of four pieces of information: (a) the combined effect of all treatments (i.e., sigma scores), (b) the unique effect of individual treatments (i.e., delta scores), (c) the overall metabolic entropy (i.e., entropy scores), and (d) time interval (relevant for time- and order-dependent combinations). To determine sigma and delta scores, we adapted a strategy previously used for creating joint chemogenomic profiles^36,37^. Specifically, we binarized drug and media flux profiles based on differential flux activity in comparison to baseline (i.e., GEM simulation without additional constraints). Sigma scores were defined as the union of binarized flux profiles for all treatments involved in a combination. Delta scores were defined as the symmetric difference between flux profiles (see ***Methods*** for details). To account for metabolic entropy, we first calculated entropy as defined by Zhu *et al*.^15^ for each drug and media flux profile. We then defined entropy scores as the mean and sum of entropy among all treatments involved in a combination. Finally, the time feature was defined as the time interval between the first and last treatments for a combination (see ***Methods*** and ***Fig. S2*** for further details).

Using feature (i.e., joint profiles) and outcome (i.e., interaction scores, IS) information for a set of drug combinations, we trained ML models to associate feature patterns to drug combination outcomes. Next, we used the trained ML models to predict outcomes for new drug combinations based on their feature information alone. We then compared our predictions against experimental data by calculating the Spearman correlation. We also assessed model performance by calculating the area under the receiver operating curve (AUROC) for both synergy and antagonism. High and positive values for both metrics indicate that model predictions correspond well with actual drug interaction outcomes.

### CARAMeL predicts drug interactions with high accuracy

We benchmarked CARAMeL against previous approaches by directly comparing our prediction accuracy against those reported in literature and those re-calculated using omics data directly instead of using flux data. For these comparisons, we trained ML models and evaluated their performance for five different cases:

1. Predicting novel pairwise drug interaction outcomes for *E. coli*^36^
2. Predicting novel three-way drug interaction outcomes for *E. coli*^37^
3. Predicting pairwise drug interaction outcomes for *E. coli* cultured in a novel nutrient condition (M9 glycerol media)^37^
4. Predicting novel pairwise and three-way interaction outcomes for *M. tb*^38^
5. Predicting interaction outcomes for multi-drug TB regimens used in clinical trials^55^

Of note, the first, second, and fourth cases tested the model’s ability to predict unseen combinations involving test drugs with new mechanisms of action. The third case assessed whether the model could predict drug interaction outcomes in a new growth environment, while the fifth case ascertained if predicted outcomes corresponded with clinical efficacy. ***Fig. 2*** summarizes our findings for all analyses listed above. For all these studies, the same train-test datasets were used for evaluating CARAMeL against the original methods to ensure direct comparison. The same thresholds for synergy and antagonism defined in the original studies were also used in all these comparisons. When re-evaluating omic-based approaches, we followed the exact procedure as reported in their respective original literature^36–38^. To ensure fair comparison between CARAMeL and omic-based approaches, we also evaluated the omic-based methods for different parameter values and report the overall best results for all datasets (***Table S1)***. Further discussion on ML model development and results, including the specific train-test allocation of interaction data reported in literature for each case, is provided below.

**Fig. 2.**
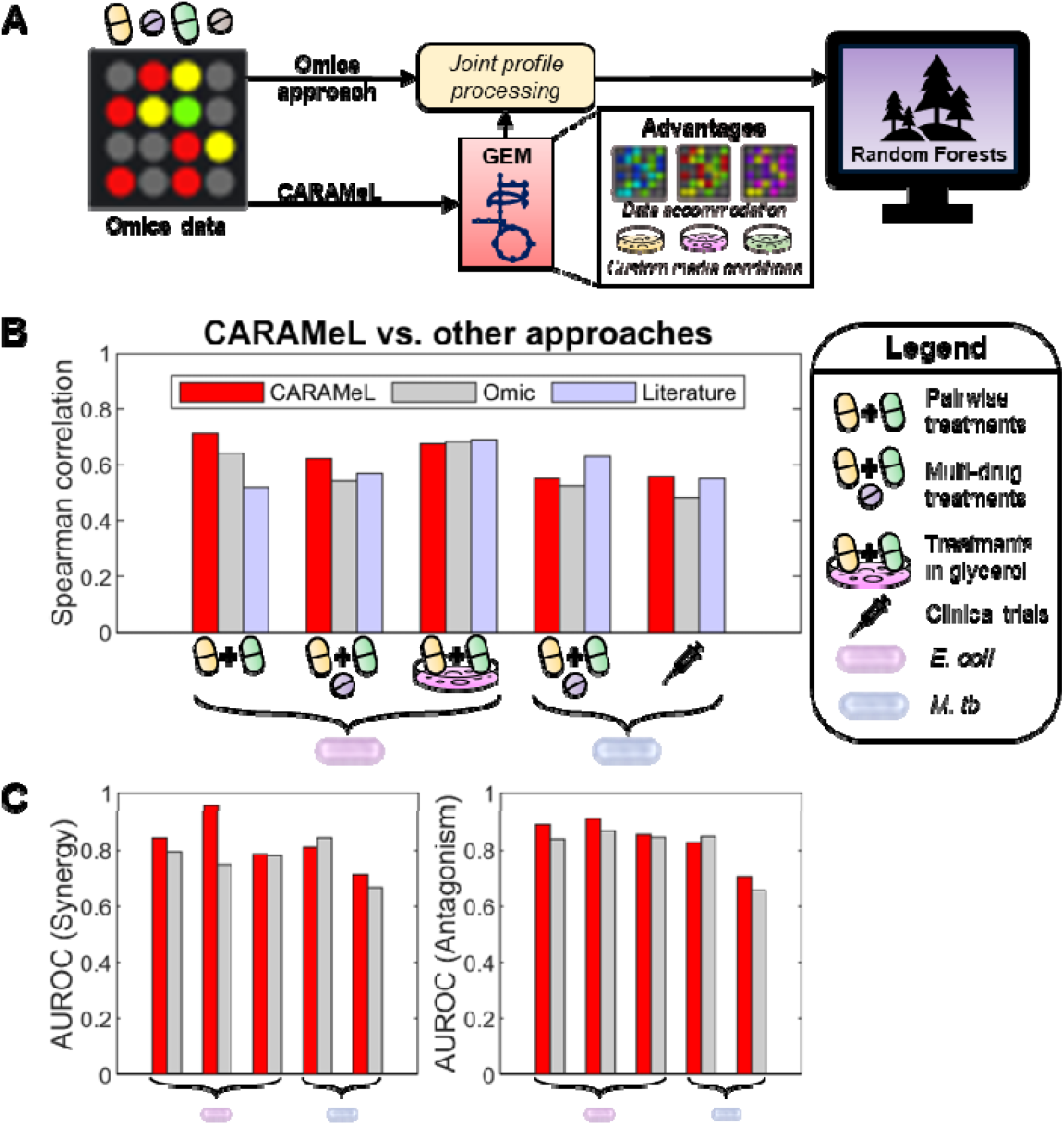
CARAMeL was benchmarked against other predictive approaches. (**A**) CARAMeL differs from previous omics-based approaches by using simulated metabolic flux data to define the joint profiles that are used in the random forests model. CARAMeL also provides advantages over the omics-based approaches, including the accommodation of diverse data types (e.g., chemogenomics, transcriptomics) and evaluation of user-defined media effects. (**B**) The Spearman correlation between actual outcomes and model predictions are shown and compared between three approaches: CARAMeL (this study), omics (determined using chemogenomic or transcriptomic data as input), and literature (reported in literature). (**C**) The area under the receiver operating curve (AUROC) for classifying interactions as synergistic or antagonistic is also directly compared between CARAMeL and omic-based approaches.

For case 1, we used drug interaction data previously measured for 171 pairwise combinations involving 19 drugs that cover a diverse set of targets^36^ (***Table S2***). Out of this total, 105 interactions involving 15 drugs were used for model training and the remaining 66 interactions, which involved four new drugs that introduced new mechanisms of action (e.g., RNA synthesis), were used for model validation. The CARAMeL model yielded significant correlations between experimental and predicted scores (R = 0.71, p ~ 10^−11^, ***Fig. S3A***). Model predictions also yielded high AUROC values for classifying synergy (IS < −0.5, AUROC = 0.84) and antagonism (IS > 2, AUROC = 0.89) (***Fig. S3B***) based on thresholds defined in the original study. Of note, these results were considerably better than those reported in literature (R = 0.52)^36^ and those re-calculated using the omic-based approach (R = 0.64) (***Fig. 2***).

For case 2, we re-trained the CARAMeL model using 171 pairwise interactions to predict 56 three-way combinations involving eight antibiotics^37^ (***Table S2***). Our model generated accurate predictions (R = 0.62, p ~ 10^−7^, ***Fig. S3C***) and notably identified synergistic interactions (IS < −0.2, AUROC = 0.95, ***Fig. S3D***) with higher accuracy than the omics-based approach (AUROC = 0.76, ***Fig. 2***).

For case 3, the CARAMeL model was once again re-trained with the 171 pairwise interactions and additional pairwise data measured for *E. coli* cultured in M9 glucose and lysogeny broth (LB) media. We then applied our model to predict 55 pairwise interaction outcomes for *E. coli* cultured in M9 glycerol media. Our model yielded results comparable to those from literature^37^ and re-determined using omics data across all three performance measures (R = 0.68, p ~ 10^−8^, ***Fig. 2, S3E, and S3F***).

For case 4, we trained a CARAMeL model using combination data for *M. tb* treated with 196 pairwise to five-way interactions involving 40 drugs^38^ (***Table S3***). We then used data for 36 unseen interactions for model validation. The CARAMeL model yielded predictions that significantly correlated with experimental data (R = 0.55, p ~ 10^−4^, ***Fig. S4A***) and performed well in classifying synergistic (IS < 0.9, AUROC = 0.81) and antagonistic (IS > 1.1, AUROC = 0.83) interactions (***Fig. S4B***). Though the CARAMeL-based correlation is slightly lower than that reported in literature^38^ (R = 0.63), our model classified both synergistic and antagonistic interactions with high accuracies that are comparable to a model trained on omics data (***Fig. 2***).

For case 5, we used the same CARAMeL model from case 4 to predict interaction outcomes for 57 multi-drug TB regimens involving nine drugs prescribed in separate clinical trials^55^ (***Table S3***). Of note, interaction outcomes for this dataset measured regimen efficacy based on sputum clearance after two months of treatment. We found that model predictions were significantly correlated (R = 0.56, p ~ 10^−6^, ***Fig. S4C***) with sputum clearance, and that model predictions classified as synergistic (IS < 0.9) captured most of the efficacious treatments (sputum clearance > 80%) amongst all 57 TB regimens (***Fig. S4D***). These results were comparable to both literature-^38^ and omic-based results across all three performance measurements (***Fig. 2***).

Overall, we found that our approach retained high accuracies in predicting combination therapy outcomes for a diverse set of test cases based on *E. coli* and *M. tb* data. This is striking considering that CARAMeL solely relies on simulated metabolic information, which was determined using only ~25–35% of available omics data.

### Using CARAMeL to predict sequential interactions

Current approaches for predicting combination therapy outcomes focus on drug treatments that are given simultaneously. Here, we extended our approach to predict treatment efficacy for time- and order-dependent (i.e., sequential) interactions. In contrast to simultaneous combinations, the order and length of each drug treatment dictates how a pathogen adapts itself, and in turn, influences its sensitivity to successive drug treatments. As such, interaction outcomes are interpreted as leading to collateral sensitivity (analogous to synergy) or cross-resistance (analogous to antagonism). For this task, we used data for *E. coli* evolved in single drug treatments over three timespans (10, 21, and 90 days) then subsequently treated with a second drug^53,56,57^. To account for both time- and order-dependent drug effect, we re-defined the delta scores for sequential joint profiles. Briefly, delta scores were defined as the difference in binarized drug profiles scaled by the time interval between treatments (mathematically defined for pairwise sequences below):

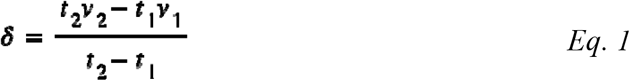

where δ = delta scores, *t* = length of treatment time, and *v* = binarized flux profile.

To initially assess how well the CARAMeL approach could predict sequential treatment outcomes, we first conducted a 10-fold cross-validation of the sequential data (N = 628), which involved 27 unique drugs (***Table S4***). We found that CARAMeL predictions moderately, but significantly, correlated with experimental outcomes (R = 0.49, p < 10^−16^, ***Fig. 3***). Further, the model performed well in determining whether a sequential interaction resulted in collateral sensitivity (IS < −0.1, AUROC = 0.73) or cross-resistance (IS > 0.1, AUROC = 0.75) (***Fig. 3***).

**Fig. 3.**
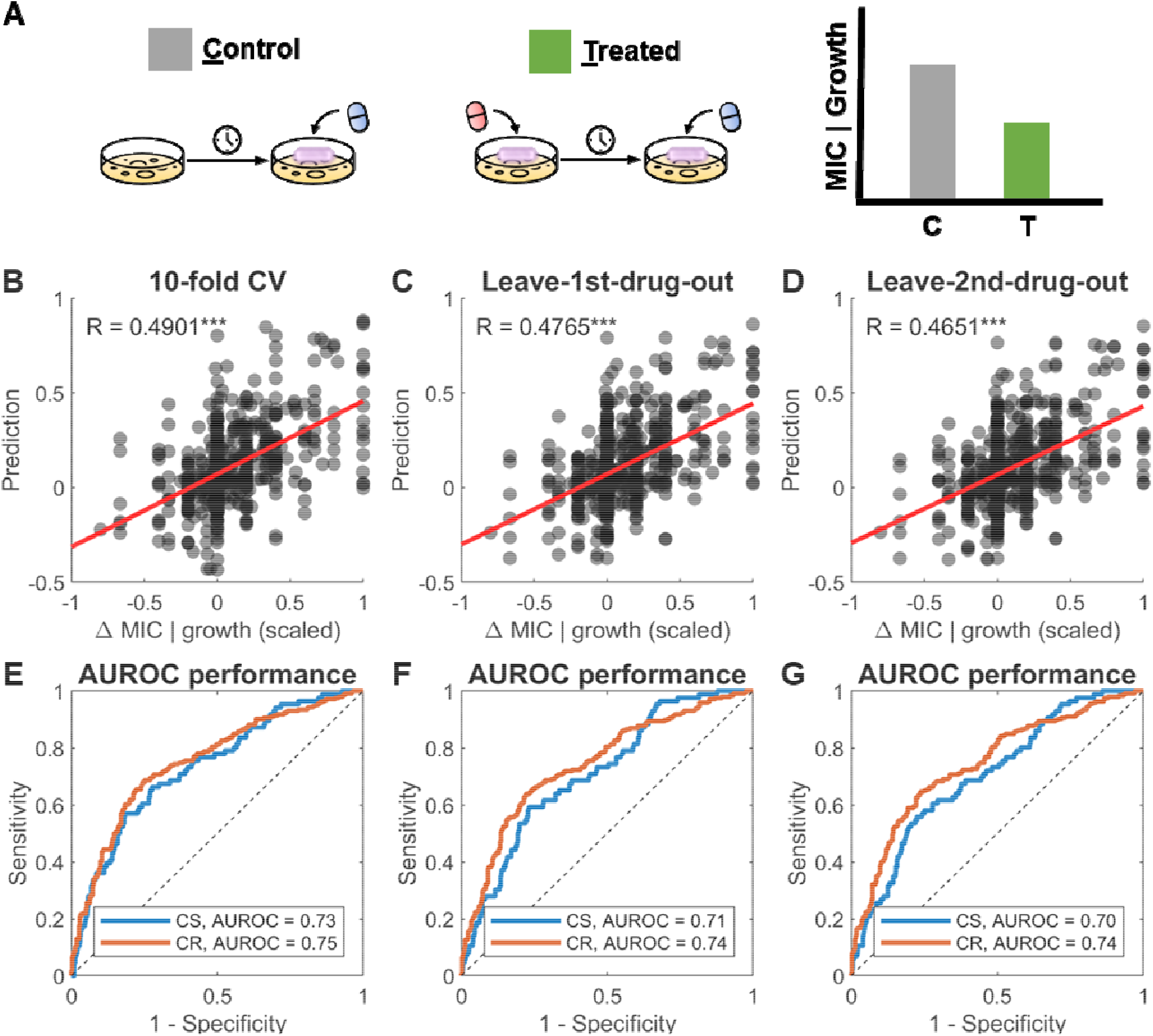
Model performance for sequential drug interactions. (**A**) The sequential treatment data used in this work measured outcomes based on the change in the minimal inhibitory concentration (MIC) or bacterial growth for a second antibiotic treatment given after exposure to a first antibiotic and compared to an untreated control (i.e., no pre-treatment with a first antibiotic). CARAMeL model performance using sequential data was evaluated based on 10-fold cross-validation (CV), leave-first-drug-out, and leave-second-drug-out analyses. Shown are the (**B-D**) scatter plots between experimentally measured outcomes (change in MIC or growth) vs. model predictions and (**E-G**) the AUROC performance for detecting collateral sensitivity (CS, outcome < 0) or cross-resistance (CR, outcome > 0). AUROC: area under the receiver operating curve. ***p-value < 10^−3^.

We next evaluated the extent of our model’s predictive power by conducting two types of leave-out analyses: (a) leave-first-drug-out and (b) leave-second-drug-out. The first case tested whether the model could generalize sequential treatment outcomes for an unknown evolved strain, while the second case assessed whether the model could generalize the immediate effect of a drug on strains evolved in other drugs. For a leave-out analysis, all interactions involving the drug of interest in the appropriate sequence position (first or second) were left out of model training and instead predicted for by the trained model. Similar to the cross-validation analysis, model performance was measured by the overall Spearman correlation and AUROC values for collateral sensitivity and cross-resistance. We found that both leave-out analyses yielded accuracies similar to those attained from cross-validation (***Fig. 3***). Overall, these results indicate that CARAMeL generates robust and accurate predictions for sequential interactions.

To gain mechanistic insight into which factors influence combination therapy outcomes, we trained a CARAMeL model using all interaction data available for *E. coli*. We then ranked features by their predictive importance based on how the model accuracy decreases when a feature is removed (see ***Methods*** for details). In total, we found that 580 features explained 95% of the variance in model predictions (***Data S1***). Of note, entropy features were amongst the top 20, implying that metabolic disarray due to antibiotic stress is indicative of treatment efficacy (***Fig. S5***). For the GEM-derived features (i.e., sigma and delta scores), we determined that the differential flux through 167 metabolic reactions associated with the top features significantly distinguished between synergistic and antagonistic interactions (two-sample t-test, adjusted p-value < 0.05, ***Data S2***). We then deduced that 8 metabolic pathways were enriched by this set of 167 reactions (hypergeometric test, adjusted p-value < 0.05, ***Table 1***). Differential activity through these pathways aligned with the expected metabolic response to antibiotic treatments. For example, increased flux through DNA repair systems (e.g., nucleotide salvage) is expected after exposure to quinolones, which target DNA gyrase^58^. Differential flux through transport reactions is also a common tactic that decreases drug concentrations within the bacterial cell, therefore minimizing their adverse effects^59^.

**Table 1.** Metabolic pathways enriched amongst top predictors for the *E. coli* CARAMeL model. Pathway enrichment was determined based on 580 features explaining 95% of the variance in model predictions. These features mapped to 333 reactions in the *E. coli* GEM iJO1366, out of which 167 had differential flux that significantly distinguished between synergy and antagonism (two-sample t-test, adjusted p-value < 0.05). Based on this 167-reaction list, 8 pathways were found to be significantly enriched (hypergeometric test, adjusted p-value < 0.05). N = number of reactions in pathway, Ratio = N / total reactions in pathway, P-value = hypergeometric test adjusted p-value.

| Pathway | N | Ratio | P-value |
| --- | --- | --- | --- |
| Purine and Pyrimidine Biosynthesis | 9 | 0.36 | 3E-05 |
| Pyruvate Metabolism | 5 | 0.50 | 1E-04 |
| Inorganic Ion Transport and Metabolism | 19 | 0.17 | 1E-04 |
| Transport, Inner Membrane | 37 | 0.11 | 1E-03 |
| Glycine and Serine Metabolism | 4 | 0.29 | 6E-03 |
| Glycolysis/Gluconeogenesis | 5 | 0.23 | 8E-03 |
| Nucleotide Salvage Pathway | 17 | 0.12 | 1E-02 |
| Cell Envelope Biosynthesis | 15 | 0.12 | 3E-02 |

### CARAMeL simulates the impact of intrinsic and extrinsic metabolic heterogeneity on drug interactions

*In vivo* metabolic conditions span growth in diverse substrates such as sugars, nucleotides, glycerol, lipids, and hypoxic conditions^44^. In contrast to existing approaches for drug combination design, CARAMeL enables drug interaction predictions in a large array of metabolic conditions. This can help prioritize drug combinations for successful clinical translation considering that the predominant nutrient source can change depending on where bacteria reside inside the host^11^. By screening different conditions that are representative of *in vivo* environments, we can identify drug combinations that target *E. coli* in diverse metabolic conditions. Moreover, evaluating drug combinations based on efficacy across a large compendium of metabolic network states will ensure robustness against heterogeneity.

To demonstrate the power of using CARAMeL in predicting condition-specific combination therapy outcomes, we applied it to predict pairwise drug interactions in multiple media conditions. For this task, we gathered experimental data for *E. coli* treated with four single drug treatments (Aztreonam (AZT), Cefoxitin (CEF), Tetracycline (TET), Tobramycin (TOB)) and two pairwise drug treatments (CEF + TET, CEF + TOB) (***Table S5***). Of note, this treatment panel evaluated the metabolic response in *E. coli* to bactericidal (i.e., death-inducing) and bacteriostatic (i.e., growth-inhibiting) drugs, both individually and in combination. Each drug treatment outcome was assessed in *E. coli* cultured in Biolog phenotype microarray (PM)^60^ plate-1, which measured metabolic respiration in 95 carbon sources and one negative control (***Fig. 4***). Out of these 95 media conditions, 57 could be simulated based on the metabolites annotated in the *E. coli* GEM (***Data S3***). As a result, ML model development and all downstream analyses were conducted using the data subset pertaining to the 57 media conditions that were simulated.

**Fig. 4.**
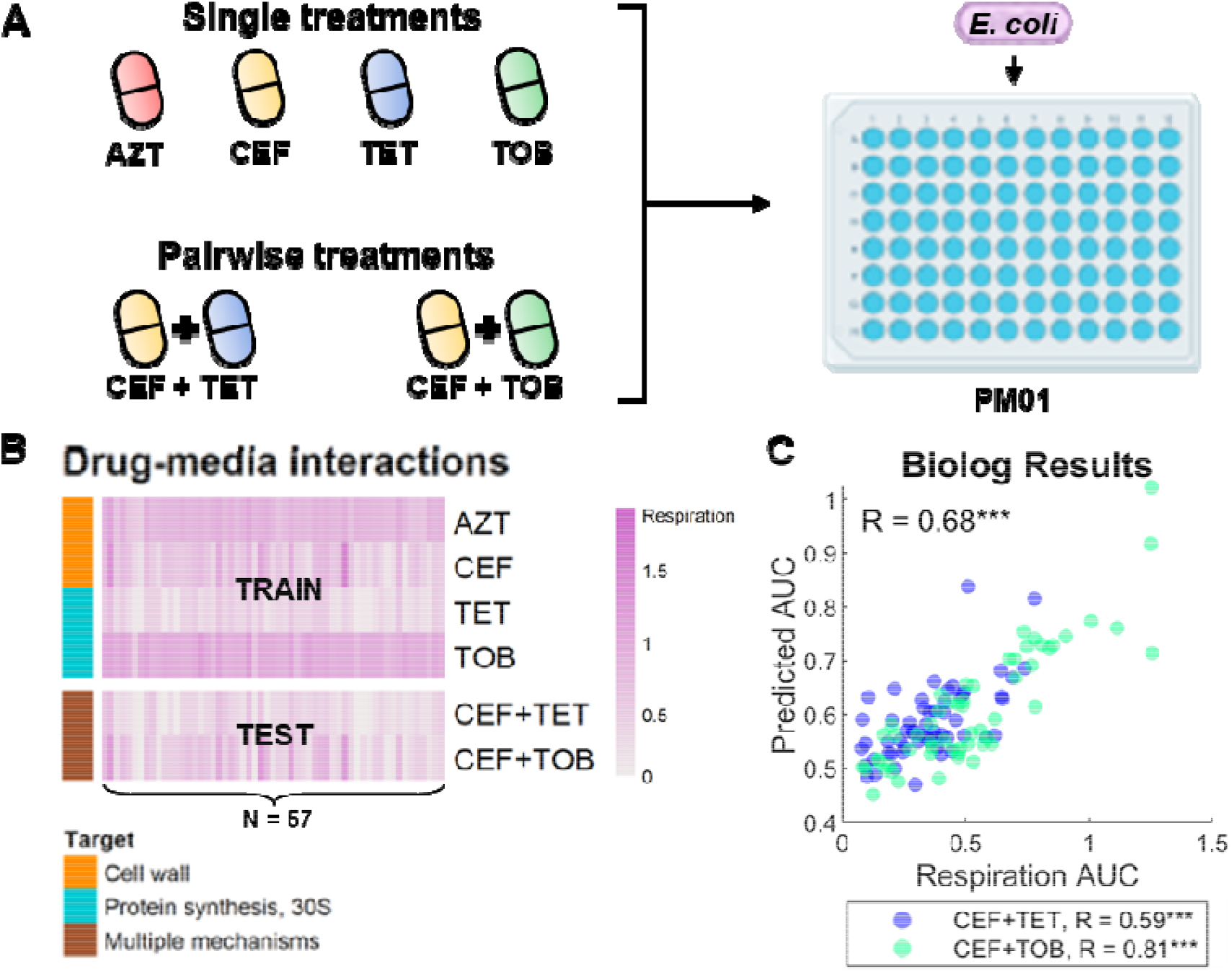
CARAMeL accurately predicted drug interaction outcomes in 57 carbon sources. (**A**) *E. coli* was cultured in 96 carbon sources (Biolog PM01 plate), then treated with four single drug treatments (AZTreonam, CEFoxitin, TETtracycline, TOBramycin) and two pairwise treatments (CEF + TET and CEF + TOB). (**B**) Heatmap of metabolic activity (measured based on the respiration ratio between treatment vs. control) in response to all experimental perturbations (data only shown for the 57 media conditions simulated using the *E. coli* GEM). (**C**) Spearman correlation between experimental outcome and model predictions for all combinations in the test set are shown. GEM: genome-scale metabolic model. *** p-value < 10^−3^.

We constructed a ML model using the following inputs: flux profiles for the four drug treatments as well as the 57 media conditions, and interaction outcomes for 228 (4 * 57) drug-media combinations. We then evaluated our model by predicting outcomes for 114 (2 * 57) drug-drug-media combinations (***Fig. 4***). Overall, we found that model predictions significantly correlated with experimental outcomes (R = 0.68, p < 10^−16^, ***Fig. 4***). We also assessed correlations specific to each drug pair and found that model predictions still corresponded well with experimental data (CEF+TET: R = 0.59, p ~ 10^−6^, CEF+TOB: R = 0.81, p < 10^−16^). This large-scale inspection of combination therapy outcome in different growth environments was only possible with the CARAMeL approach, where flux profiles could be determined for 57 media conditions. A direct comparison of the same scale was not possible with the omic-based approach, as neither chemogenomic nor transcriptomic data was available for all these media conditions.

We next evaluated how cell-to-cell heterogeneity influenced combination therapy outcomes using population FBA^61^, a modeling approach that simulates cell-specific metabolic heterogeneity based on single-cell proteomics data^62^. Specifically, information on the protein copy number levels measured for *E. coli* cultured in M9 glucose media is used to constrain the metabolic model^62^. To simulate heterogeneity between cells, the single-cell proteomics data is randomly sampled based on the Gamma distribution for each cell and subsequently used to constrain the GEM to simulate cell-specific metabolic states (***Fig. 5***). We used population FBA to simulate 1,000 *E. coli* cells cultured in M9 glucose media (***Methods***). We then generated pairwise predictions between all drugs for which the *E. coli* CARAMeL model was trained on (N = 33) against the 1,000 simulated cells (***Data S4***). For each drug pair, we evaluated three interaction cases: (a) simultaneous treatment (D_1_ + D_2_), (b) sequential treatment from D_1_ to D_2_, and (c) sequential treatment from D_2_ to D_1_. For sequential interactions, we set the duration for the first treatment to 14 days, based on the most commonly prescribed antibiotic treatment duration against bloodstream infection by *Enterobacteriaceae*^63^, and one day for the second treatment.

**Fig. 5.**
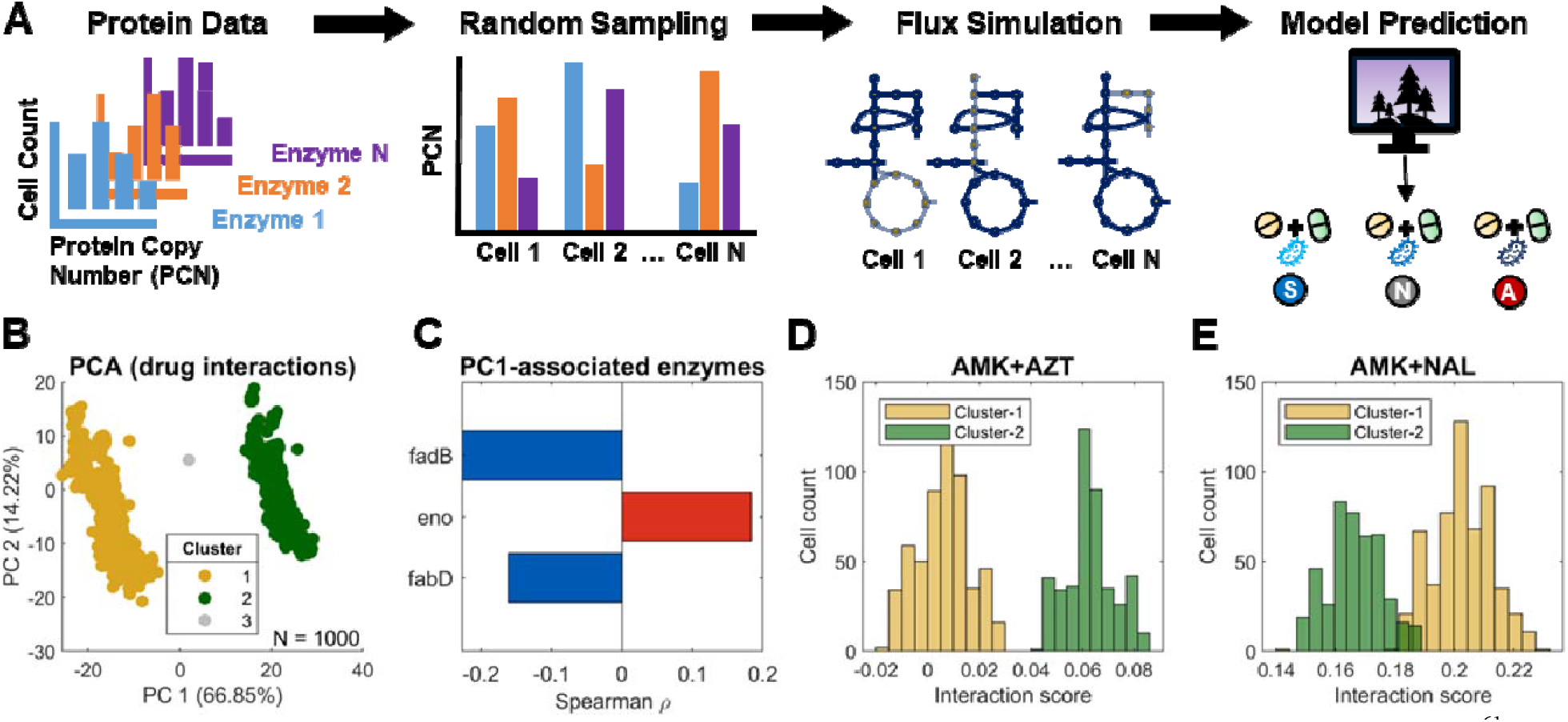
Single cell-specific combination therapy predictions. (**A**) Schematic showing how population FBA^61^ was applied to simulate cell-specific metabolic heterogeneity. Specifically, single-cell proteomics data^62^ was randomly sampled (based on the Gamma distribution) and used to constrain the *E. coli* genome-scale metabolic model to simulate cell-specific fluxes, which were ultimately used to generate cell-specific drug interaction predictions. (**B**) Cells were found to cluster into two distinct groups after applying principal component analysis (PCA) onto the simultaneous prediction data (PC loadings for the first two dimensions reported in ***Data S6***). (**C**) The sampled level for three enzymes (eno, fadB, and fabD) were found to significantly correlate with cell-specific scores along principal component 1 (PC1) from panel B. (**D**) Cluster-1 cells were predicted to be more sensitive to most drug combinations (e.g., AMK + AZT) compared to cluster-2 cells. (**E**) For a smaller set of drug combinations (15%), primarily involving quinolones (e.g., AMK + NAL), cluster-2 cells were predicted to be more sensitive than cluster-1 cells. AMK: amikacin, AZT: aztreonam, NAL: nalidixic acid.

Using the prediction landscape for the 1,000 cells, we determined the extent of cell-to-cell variability for each unique drug pair (***Data S5***). Overall, sequential predictions varied more largely between cells (up to 14% change in standard deviation relative to the mean) while there was less than 5% change in standard deviation compared to the mean for the simultaneous case, suggesting that simultaneous treatments may be more robust to heterogeneity (***Fig. S6***). Interestingly, the variation among simultaneous predictions tended to follow a bimodal distribution (***Fig. S7***). Applying principal component analysis (PCA) onto the simultaneous prediction landscape for all 1,000 cells confirmed that two distinct sub-populations can be determined via k-means clustering (***Fig. 5 and S8***, see ***Methods*** for details). We further confirmed that the cell clustering was not being driven by non-uniform sampling of the flux solution space by re-sampling single-cell fluxes using optGpSampler^64^ (***Fig. S9***). Of note, this distinct cell grouping was not observed when PCA was applied to cell-specific proteomics data nor metabolic flux data (***Fig. S10***).

We next sought to determine whether the clustering pattern seen in the drug interaction PCA plot was being driven by specific enzyme levels or metabolic activity. Thus, we evaluated the correlation between the cell-specific scores along principal component 1 (PC1) against corresponding enzyme levels and simulated flux profiles. We determined that sampled levels for three enzymes (eno, fadB, and fabD) significantly correlated with the cell mapping along PC1 (***Fig. 5***). These enzymes correspond to enolase (involved in glycolysis), a multi-functional enoyl-CoA hydratase (involved in lipid metabolism), and malonyl-CoA-acyl carrier protein transacylase (involved in lipid metabolism), respectively. A similar comparison of PC1 scores with the simulated flux data revealed more than 400 significantly associated reactions (***Data S7***), which altogether correspond to 16 pathways (hypergeometric test, adjusted p-value < 0.05, ***Fig. S11***). These findings confirm that cluster-1 cells differ from cluster-2 cells based on their metabolic activity through glycolysis (eno) and lipid metabolism (fadB and fabD).

Hence, fluctuations in the levels of these three proteins were predicted to drive the broad metabolic shift between the two sub-populations. To confirm the causality, we performed knockouts of these three proteins. A similar PCA assessment for cells simulated to have single- and multi-gene knockout of eno, fadB, and fabD confirmed that the PCA-based cell clustering seen in ***Fig. 5*** is strongly driven by the metabolic states characterized by eno and fabD levels (***Fig. S12***).

Based on the direction of the enzyme correlation with PC1 scores, we inferred that cluster-1 cells exhibit low glycolysis and high lipid metabolism, while cluster-2 cells exhibit the opposite behavior. For a large set of drug combinations (85%), we found that reduced glycolysis coupled with high lipid metabolism promoted more synergistic outcomes (***Fig. 5, Data S8***), while the same metabolic state was found to promote more antagonistic outcomes for a smaller set (15% of the combinations) (***Fig. 5, Data S8***). Of note, a small number of cells representing <1% of the total population (labeled as “cluster-3”) did not cluster together with either dominant sub-population but instead fell near the center of the drug interaction PCA space (***Fig. 5 and S7***). Closer inspection of enzyme levels for eno, fadB, and fabD revealed that the sampled levels for eno were much lower for cluster-3 cells (~400 per cell) compared to cluster-1 and cluster-2 cells (~600 per cell, ***Fig. S13***). Considering that enolase is an essential enzyme for maintaining glycolysis and cell growth, an adequately high level for enolase (e.g., > 500 per cell) may be required to simulate stable flux solutions that lead to the patterns we observe in ***Fig. 5***. These cells may represent an unstable transition state between cluster 1 and 2.

Interestingly, the smaller set of drug combinations with antagonistic outcomes in cluster 1 is overrepresented by combinations that include quinolones such as nalidixic acid (~12% among synergistic vs. ~86% of antagonistic combinations in cluster-1 involved quinolones, refer to ***Data S8***). A prior study has found that quinolone efficacy is reduced in high-density bacterial populations, likely due to depletion of metabolites that couple carbon metabolism to oxidative phosphorylation^65^. The same study subsequently shows that quinolone efficacy can be restored via supplementation with glucose and an electron acceptor, which stimulate respiratory metabolism. Our findings, coupled with the literary evidence described above, indicate that cluster-1 cells may represent a sub-population that is tolerant to treatments involving quinolones. Given the highly interconnected nature of cellular metabolism, stochastic changes in a small number of key metabolic enzymes can result in distinct phenotypes when treated with stressors. These two sub-populations may not be evident in an unperturbed system which shows fluctuations in numerous proteins; however, when exposed to antibiotics they may result in bifurcation into two stable sub-populations. Though experimental validation would be required to fortify these results, we confirmed that all the reported findings tied to the cell-specific predictions are robust to a wide range of modeling parameters (***Fig. S8***).

### Screening for robust combination therapies

Synergy observed in the lab may not result in synergy *in vivo* due to differences in growth conditions or drug pharmacokinetics, wherein drugs may reach the infection site at different times rather than simultaneously^66^. Considering these factors, combination therapies that show synergy across growth conditions and time scales hold the best potential for successful clinical translation. To discover such therapies, we predicted pairwise and three-way regimen outcomes for all drugs for which the *E. coli* CARAMeL model was trained on (N = 33) across 57 carbon sources (from Biolog PM01). For sequential interaction predictions, treatment duration for pairwise treatments was set to 14 days followed by 1 day, while three-way treatments were set to a 14-14-1-day prescription. In total, we generated predictions for 90,288 pairwise combinations (_33_C_2_ pairs × 3 interaction cases × 57 PM01 conditions, ***Data S9***) and 2,176,944 three-way combinations (_33_C_3_ combinations × 7 interaction cases × 57 PM01 conditions, ***Data S10***). Out of 528 unique drug pairs and 5,456 unique three-way interactions, none was predicted to be synergistic across all media conditions and interaction cases. In fact, sustained synergy across media conditions seems to occur for only a small subset (< 10%) of drug interactions (***Fig. S14***). Specifically, 73 drug pairs and 165 three-way interactions were predicted to yield synergy both simultaneously and sequentially in at least one media condition (***Fig. 6***). Of note, all 73 drug pairs showed less than 5% cell-to-cell variation based on population FBA for all interactions cases (i.e., simultaneous and sequential interactions). Upon closer inspection of both pairwise and three-way sets, synergy was not found to be retained across a majority of media conditions for three-way drug interactions. On the other hand, several pairwise interactions were found to retain synergy well; specifically, 24 drug pairs out of 73 were found to be synergistic in more than 50% of conditions in both simultaneous and sequential cases.

**Fig. 6.**
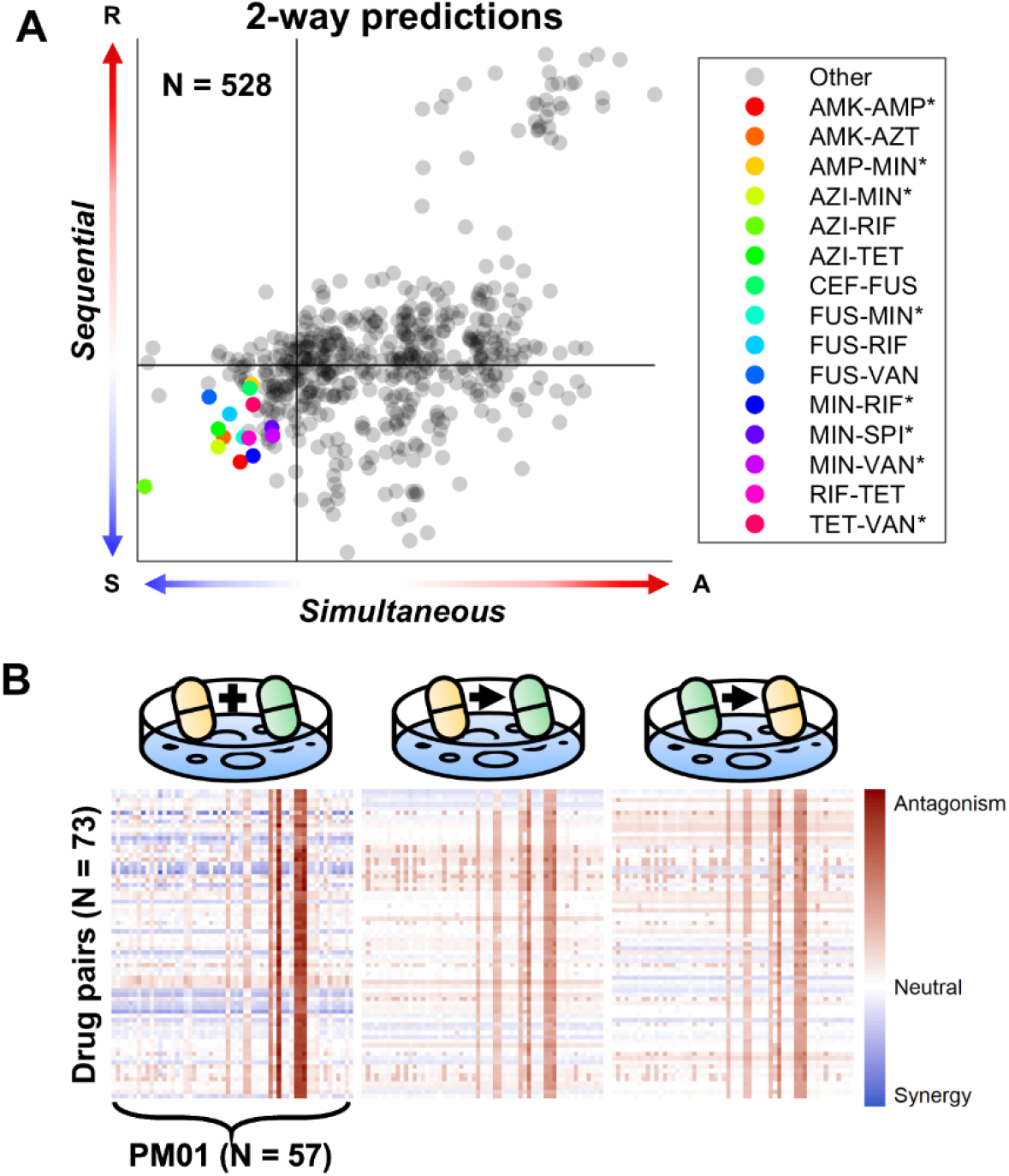
Pairwise combination therapy prediction across 57 media conditions. (**A**) Out of 528 drug pairs, 73 were predicted to yield synergy (IS < 0) in at least one media condition for both the simultaneous and sequential cases (the top 15 robustly synergistic drug pairs are listed in the legend). (**B**) Heatmap of the predicted interaction scores for the 73 drug pairs across 57 media conditions and three interaction types (D_1_ + D_2_, D_1_ → D_2_, D_2_ → D_1_). Refer to ***Tables S2 and S4*** for full descriptions on antibiotics used for *E. coli*.

Interestingly, several of these 24 drug pairs possess evidence for clinical use against bacterial infections. For example, amikacin-ampicillin treatment (AMK-AMP) has previously been shown to be clinically effective for a wide range of infections^67–69^ including treatment of bacteremia in neutropenic patients^70^ and neonatal bacterial infections^71^. Other drug interactions of note include: azithromycin-rifampicin (AZI-RIF), which has demonstrated clinical efficacy in treating arthritis induced by pathogenic Chlamydia (Gram-negative)^72^; fusidic acid-rifampicin (FUS-RIF), which has shown clinical efficacy against prosthetic joint infection caused by drug-resistant staphylococci (Gram-positive)^73^; and minocycline-rifampicin (MIN-RIF), which has been shown to prevent colonization by slime-producing staphylococci in catheters^74^. Additionally, rifampicin combined with other drugs has been advised as treatment for Gram-negative and non-mycobacterial infections^42,75^. Further investigation into the set of 24 drug pairs predicted to yield robust synergy may lead to the discovery of new combination therapies that could be put to clinical use.

## Discussion

Here we introduced CARAMeL, a modeling approach to design condition-specific antibiotic regimens. CARAMeL offers multiple advantages over prior methods of similar nature. First, our approach enables use of diverse data types (e.g., chemogenomics, transcriptomics) individually or in combination, derived from a single source or in combination from multiple sources, therefore maximizing the number of drugs that are screened. For instance, our *E. coli* CARAMeL model leveraged use of both chemogenomics data (for defining drug flux profiles) and proteomics data (for simulating single cells), a feat that cannot be accomplished with prior methods of similar nature. Second, we extended our approach to simulate different interaction cases (simultaneous vs. sequential) when designing combination therapies. To our knowledge, no framework currently exists to incorporate these factors into drug discovery efforts against AMR. Third, the use of GEMs enables simulation of highly tunable metabolic conditions (as showcased with our analysis of the Biolog PM01 data), which may be leveraged to investigate combination therapy outcomes in the host environment. GEMs also enable the simulation of pathogen metabolic heterogeneity, due to both intrinsic stochasticity and the metabolic environment. Pathogen heterogeneity is a critical barrier in designing effective antibiotic therapies, and this must be factored into combination therapy design to mitigate the rise of resistant strains.

Additional advantages to the CARAMeL approach include its mechanistic model interpretability, its ability to simulate pathogen metabolic heterogeneity, and its use in generating predictions across numerous conditions in large-scale. Regarding model interpretation, the *E. coli* CARAMeL model revealed that entropy, or metabolic disarray, plays an important role in combination therapy outcome. The direct link between model features (i.e., sigma and delta scores) and GEM reactions also pinpointed pathways that are activate in response to drug treatment (***Table 1***), many of which align with the expected resistance mechanisms against antimicrobial stress^58,59^.

Using population FBA^61^, we investigated how drug interaction outcomes may differ from cell-to-cell. Our findings potentially point to a connection between the metabolic state of a cell and its tolerance against combination treatments. Specifically, we found that sensitivity to a broad-range of drug combinations may be influenced by the variation in activity of glycolysis and lipid metabolism; these processes are directly related to antibiotic action and interaction such as uptake, respiration, and oxidative stress^76^. Our results also imply that drug interaction outcomes measured for a bulk cell population may not be representative for cell sub-populations. Surprisingly, very few cells show the “average” behavior of the population; in most cases, the average prediction may be defined by the outcome in two dominant sub-populations where one is more sensitive to treatment while the other is more tolerant. This investigation, along with our results with the Biolog data, demonstrate how pathogen metabolic heterogeneity may arise due to both intrinsic stochasticity and the local growth environment. Pathogen heterogeneity is a critical barrier in designing effective antibiotic therapies, and this must be factored into combination therapy design to mitigate the rise of resistant strains.

Finally, analysis of the drug interaction landscape suggests that only a small set of 24 out of ~6,000 combinations show robust synergy across growth conditions and interaction cases, with some possessing clinical evidence for efficacy^42,68–75^. Further investigation into this list of drug interactions may lead to the discovery of new combination therapies for clinical application.

Ultimately, CARAMeL serves as a proof-of-concept of how computational approaches, such as systems-level metabolic modeling and machine learning, can be combined to create hybrid models that provide mechanistic insight into various biological processes^77–79^, in this case antimicrobial efficacy and resistance. Although the use of GEMs in CARAMeL offers major advantages with data compatibility, condition tunability, and mechanistic insight, it also introduces some limitations. The level of accuracy and thoroughness in GEM annotation may influence CARAMeL model performance. Moreover, our current approach only provides a “snapshot” perspective of the metabolic response to a condition. This may be a potential reason for the diminished CARAMeL model performance in predicting sequential outcomes. Nevertheless, these are areas that can be addressed with continued curation of GEMs^80^ and advances in dynamic metabolic modeling^81,82^. Overall, the ability to simulate specific growth environments and pathogen metabolic heterogeneity offers the potential to evaluate treatment efficacy *in vivo* and advance clinical translation of novel antibiotic regimens. Moreover, these combination therapies could restore use of defunct antibiotics against resistant pathogens while mitigating further resistance^24,83^. Beyond bacterial infections, CARAMeL has the potential to design explainable combination therapies that are urgently needed to combat fungal infections^84^ and drug-resistant cancer cells^85^. Such broader applications can be achieved by leveraging the large volume and diversity of highly curated GEMs that exist and continue to be constructed^86^. Our approach can further be used to understand the role of metabolic heterogeneity in cancer treatment, which plays a major role in tumor drug resistance^87,88^.

## Methods

### Experimental Design (Biolog Phenotype Microarray)

*E. coli* MG1655 was cultured in Biolog phenotype microarray (PM) 1, which screened bacterial growth in 95 carbon sources and a negative control (i.e., water)^60^. *E. coli* was subsequently treated with six distinct drug treatments in duplicate: aztreonam (0.03 ug/mL), cefoxitin (1.87 ug/mL), tetracycline (1.42 ug/mL), tobramycin (0.15 ug/mL), cefoxitin (1.87 ug/mL) + tetracycline (1.42 ug/mL), and cefoxitin (1.87 ug/mL) + tobramycin (1.42 ug/mL). Including a reference plate (*E. coli* growth in PM01 only), phenotype in each treatment was colorimetrically measured in duplicate using tetrazolium violet dye, which quantifies cellular respiration. All experimental procedures, data collection, and quality control were performed at Biolog, Inc. The area under the respiration curve was calculated using MATLAB and reported as the ratio of treatment to reference.

### Simulating Metabolic Flux using GEMs

The *E. coli* GEM iJO1366^48^ and the *M. tb* GEM iEK1008^49^ were used to simulate metabolic fluxes at steady-state. To simulate drug flux profiles, chemogenomic data for *E. coli*^50^ and transcriptomic data for *M. tb*^38^ served as GEM constraints. Specifically, differential gene regulation in response to each drug treatment was inferred from each dataset. For chemogenomic data, which measured single-gene knockout (KO) fitness, genes KOs that promoted growth were assumed as dispensable while gene KOs that resulted in low fitness were assumed to be essential for growth in said condition. Based on these assumptions, genes corresponding with low (z < −2) and high (z > 2) fitness were inferred to be up- and down-regulated, respectively. Of note, the original source for the *E. coli* chemogenomic data used in this study was already processed and normalized into z-scores that accounted for both the gene KO and drug treatment effects. For transcriptomic data, which measured single-gene expression, up- and down-regulation were directly inferred based on high (z > 2) and low (z < −2) expression values, respectively. These processes yielded individual sets of differentially regulated genes that were integrated into corresponding GEMs using a linear optimization version of the integrative metabolic analysis tool (iMAT)^89,90^. To determine media flux profiles, metabolite availability was computationally defined by constraining exchange reactions annotated in iJO1366. For each carbon source of interest (e.g., glycerol), the lower bound (i.e., uptake rate) for the corresponding exchange reaction (e.g., glycerol exchange) was set to −10 g/mmol to allow cellular intake.

Of note, use of the linear iMAT algorithm required constraint-based modeling (CBM) parameter fine-tuning for three variables: kappa, rho, and epsilon^91^. Kappa and rho serve as relative weights for “off” and “on” reactions associated with the differentially expressed genes, respectively, in their contribution to the objective function. Epsilon represents the minimum flux through “on” reactions. For the purposes of this research, we varied all three parameter values from 10^−3^ to 1 and determined the optimal parameter set based on three criteria: (1) maximizing the Spearman correlation between predicted and actual interaction scores after 10-fold cross-validation using a training dataset, (2) minimizing the number of conditions simulated to have no growth, and (3) ensuring non-zero variability in the simulated growth rates between conditions. ***Table S6*** provides results for all three assessments for all parameter sets of interest. The following optimal parameter values were obtained for each GEM using the training dataset: (1) iJO1366 – kappa =10^−2^, rho = 10^−2^, epsilon = 1, and (2) iEK1008 – kappa = 10^−2^, rho = 10^−2^, epsilon = 10^−2^. These parameter values were used for all results when benchmarking CARAMeL against previous approaches based on *E. coli* and *M. tb* drug interaction datasets (***Table S7***).

To simulate cell-specific flux data, we applied population FBA^61^, an approach that models metabolic heterogeneity within a cell population using proteomics data. For the purposes of this study, we defined a population of 1,000 *E. coli* cells to simulate using the default parameters for population FBA. For the reproducibility analysis (related to ***Fig. S8***), we ran population FBA for 1,000 cells and subsequently used the simulated flux data to generate cell-specific drug interaction outcome predictions a total of 30 times. To retrieve uniform sampling of the cell-specific flux solution space (related to ***Fig. S9***), we applied optGpSampler^64^ to generate 100 flux solution samples for 10 unique cells derived from a population FBA simulation. See the populationFBA.mlx file in the GitHub repository for this study (https://github.com/sriram-lab/CARAMeL) for details on the exact implementation of population FBA and optGpSampler. Of note, the flux data for all conditions (i.e., drug, media, single-cell) used to define joint profiles was generated from a single run of the condition-appropriate CBM method (i.e., iMAT, FBA, population FBA). The exact flux data that was used to generate all results is stored as a data file in the GitHub repository associated with this study (https://github.com/sriram-lab/CARAMeL).

### Data Processing to Determine Joint Profiles

Flux profiles were used to define joint profiles for each drug combination, which were comprised of four pieces of information: sigma scores, delta scores, cumulative entropy, and treatment time interval (***Fig. S2***). Sigma and delta scores were representative of the combined and unique effect of drugs involved in a combination, respectively. Of note, joint profiles for the original omics-based approaches were only defined by sigma and delta scores^36–38^. For CARAMeL, the general procedure for determining sigma and delta scores was retained from the original literature, with the input data (flux profiles) being the only difference. Both score types were determined after flux profiles were binarized based on differential flux activity (either positive or negative) in comparison to baseline, mathematically defined below:

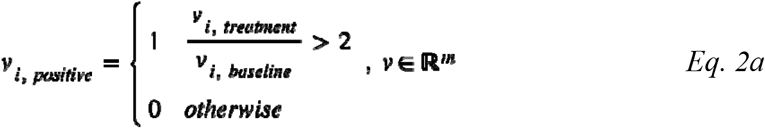

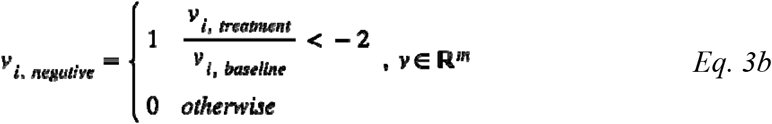

where *v* = reaction flux and *m* = total number of GEM reactions. Sigma scores were mathematically defined for both simultaneous and sequential interactions using the following equation:

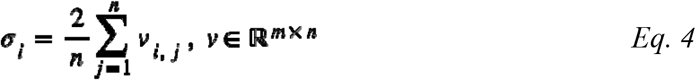

where σ = sigma score, *v* = binarized flux profile, *m* = total number of GEM reactions, and *n* = total number of conditions in a combination. Delta scores were separately defined for simultaneous and sequential interactions based on *Eq. 5* and *Eq. 5*, respectively:

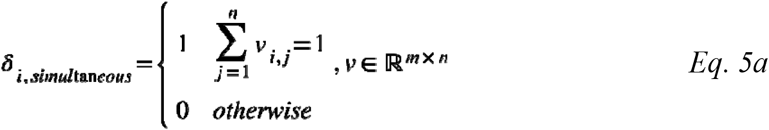

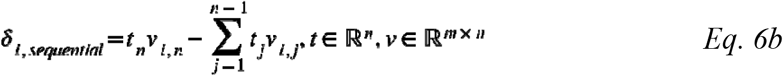

where δ = delta score, *t* = treatment time interval, *v* = binarized flux profile, *m* = total number of GEM reactions, and *n* = total number of conditions in a combination. Cumulative entropy features were determined by processing non-binarized flux profiles in two steps. First, metabolic entropy for each condition was mathematically defined by the following equation:

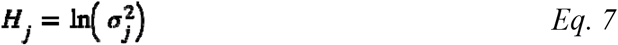

where *H*_*j*_ = metabolic entropy due to condition *j* and = variance in the non-binarized flux profile for condition *j*. Of note, this formulation was adapted from Zhu *et al*., who quantified entropy of the bacterial stress response to antibiotics^15^. Next, the mean and sum in entropy for all conditions involved in an interaction were used to define two distinct entropy features. Finally, the time feature was defined as the time interval between the first and last treatment for a combination. For simultaneous interactions, the time feature was set to zero.

### ML Model Development using Random Forests

All CARAMeL models were built in MATLAB (Mathworks, Inc.) using the regression-based Random Forests (RF) algorithm^92^. Briefly, RF is an ensemble method comprised of decision trees that learn to associate feature information to a target variable. For the regression-based approach, the RF model returns the mean prediction from all decision trees. To develop CARAMeL models, joint profiles served as feature information while drug interaction scores were used as the target variable. Interaction scores were quantified using the Loewe additivity model^93^, which is based on drug concentrations (refer to the original sources of drug interaction datasets for further details in score calculation). Both joint profiles and interactions scores for drug combinations of interest were used as model inputs during training, while only joint profiles were provided as input during model testing. Default values for all other model parameters were used during both training and testing.

### ML Model Performance Assessment

Model performance was evaluated based on two metrics: (1) the Spearman correlation between actual and predicted interaction scores and (2) the area under the receiver operating curve (AUROC) for classifying interactions as synergistic or antagonistic. Of note, model predictions for TB regimens used in clinical trials were negative transformed before being compared to clinical outcomes. Since these clinical trials reported percentage of patients that were cured, we would expect to see a negative correlation between interaction scores and clinical efficacy, with synergistic regimens (negative IS) performing better than antagonistic regimens. The sign for the scores were hence flipped to maintain a positive correlation indicating good model performance. Classification of simultaneous drug interactions was based on score threshold values reported in the original literature for a dataset. For both sequential interactions and the CARAMeL model trained on all interaction data for *E. coli*, interaction scores were first scaled by the maximum absolute value (*Error! Reference source not found*.). Interaction values below −0.1 and above 0.1 were then used to classify interactions as synergistic and antagonistic, respectively. For the 10-fold cross-validation analysis conducted for sequential interactions, the interaction data was randomly partitioned into ten subsets of similar size (N ~ 63). CARAMeL was then applied to predict each subset at a time, where the given subset was left out of the model training (i.e., the remaining 90% of the data was used to train the model). All model predictions were then compared to the sequential data as a whole to calculate the overall Spearman correlation and AUROC values.

### CARAMeL Top Feature Extraction

Top features were determined based on their ranked importance in generating accurate predictions. To calculate feature importance, each feature was first left out of model training and testing. The mean squared error (MSE) between predicted and true interaction scores was then calculated for each model. Finally, feature importance was measured as the increase in MSE for a model lacking a feature compared to the model trained on all features. After ranking features according to decreasing importance, the first set of features amounting to a cumulative importance of 0.95 (corresponding to 95% variance explained) were selected for downstream model interpretation and analysis. To determine overall importance, we trained a CARAMeL model using all interaction data available for *E. coli*. Broadly, this included three sets of simultaneous combinations^36,37^ (pairwise, three-way, and media-specific treatments) and three sets of sequential interactions^53,56,57^ (differing based on elapsed treatment time). To account for differing units of measurement between datasets, we scaled interaction scores according to the following formula:

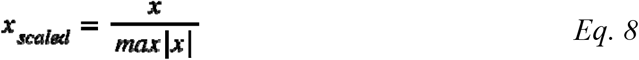

where *x* is a vector of interaction scores for a given dataset. This scaling constrained all interaction scores to range between ±1 while retaining the sign consensus for classifying interactions based on their score (negative IS → synergy, positive IS → antagonism). In total, we trained our model on 966 drug interactions and attained highly accurate predictions (R = 0.45, p < 10^−16^) for both synergistic (IS < −0.1, AUROC = 0.67) and antagonistic (IS > 0.1, AUROC = 0.71) interactions based on a 10-fold cross-validation.

### Principal component analysis (PCA) and k-means clustering to determine cell sub-populations

Principal component analysis (PCA) was applied onto three datasets pertaining to single-cell results: (1) the sampled enzyme levels (352 proteins × 1,000 cells), (2) the simulated metabolic reaction fluxes (2,583 reactions × 1,000 cells), and (3) the CARAMeL predictions for simultaneous drug interactions (528 drug pairs × 1,000 cells). PCA results were then visualized onto the 2-dimensional space defined by the first two principal components (PCs) for all three PCA applications. The cell sub-populations reported in the main text were determined via k-means clustering (k = 2) of the PCA data for drug interactions. The silhouette value (i.e., measure for evaluating cluster assignment) for each cell was subsequently calculated to determine the presence of any “outlier” cells (i.e., cluster-3, silhouette value < 0.5).

### Statistical Analysis

A one-way analysis of variance (ANOVA) test was used to compare both the entropy mean and entropy sum of drug interactions grouped by their classification (synergy, neutral, antagonism). A multiple comparison test based on Tukey’s honestly significant difference (HSD) was subsequently performed to identify statistically significant pairwise differences using a p-value threshold of 0.05. A two-sample Student’s t-test with unequal variance was used to define which reactions distinguished between synergistic and antagonistic interactions based on differential flux activity. Lastly, a hypergeometric test was conducted to determine significantly enriched metabolic pathways based on GEM reactions associated with top CARAMeL predictors. For this test, the total number of reactions annotated in iJO1366 corresponded with the population size. Of note, the p-values determined from t-tests and hypergeometric tests were adjusted using the Benjamini-Hochberg approach^94^.

## Acknowledgments

We thank Brendan Lewis at Biolog for carrying out the Phenotype Microarray assays.

## Funding

National Institutes of Health grant R35 GM13779501 (SC) National Institutes of Health NIAID R56AI150826 (SC) University of Michigan faculty start-up fund (SC)

## Competing interests

The authors declare that they have no competing interests.

## Data and materials availability

All datasets and code used within this work are provided through the CARAMeL GitHub repository (https://github.com/sriram-lab/CARAMeL). Additional information on datasets and results are available in the supplementary materials. The following software tools were used in this study: MATLAB R2021b (for analysis and visualization) and R v4.1.1 (for visualization).

## Notes

https://github.com/sriram-lab/CARAMeL

https://www.dropbox.com/s/859ebsx1drri1xc/data.xlsx?dl=0

https://www.dropbox.com/s/lmd3ejpms4095xw/CARAMeL_workspace.mat?dl=0

### Competing Interest Statement

The authors have declared no competing interest.

### Summary of Updates

New results on cell-specific drug interaction outcomes were reviewed and updated. All materials (figures, tables, data, code) were updated to reflect the aforementioned change.

