## Supplementary Materials for "A flux-based machine learning model to simulate the impact of pathogen metabolic heterogeneity on drug interactions"

**This PDF file includes:**

Figs. S1 to S15

Tables S1 to S7

Data S1 to S10

**Other Supplementary Materials for this manuscript include the following:**

Data S1 to S10 (Available through Dropbox at <https://www.dropbox.com/s/859ebsx1drri1xc/data.xlsx?dl=0>)

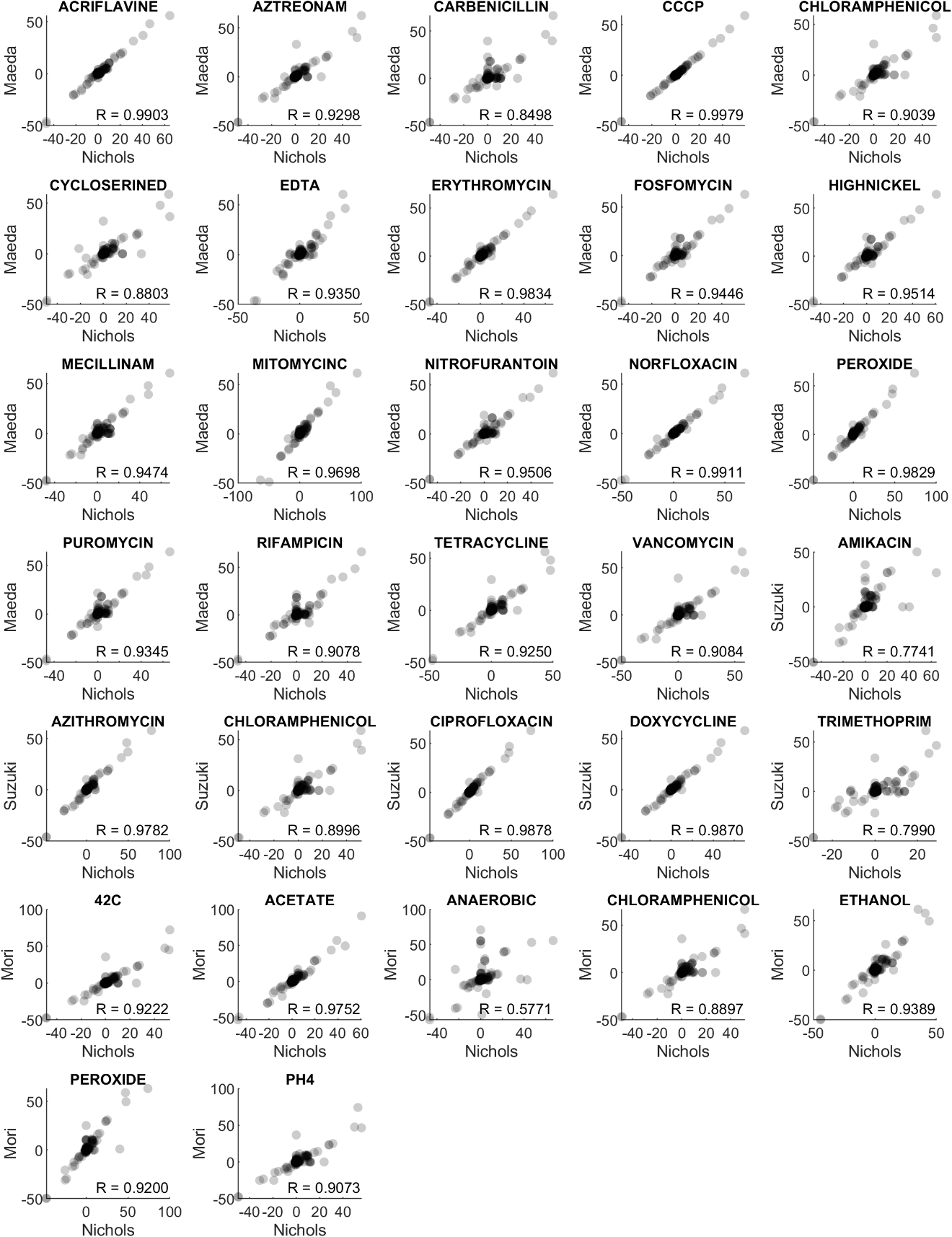

Fig. S1.

**Flux profile comparison between different omic-based simulations.** Correlations based on Pearson’s method (all yielded p << 10^-3^). All plots possess the same number of points (i.e., reactions, N = 2583). Nichols: chemogenomic-based (*50*), Maeda: transcriptomic-based (*52*), Suzuki: transcriptomic-based (*53*), Mori: proteomic-based (*54*).

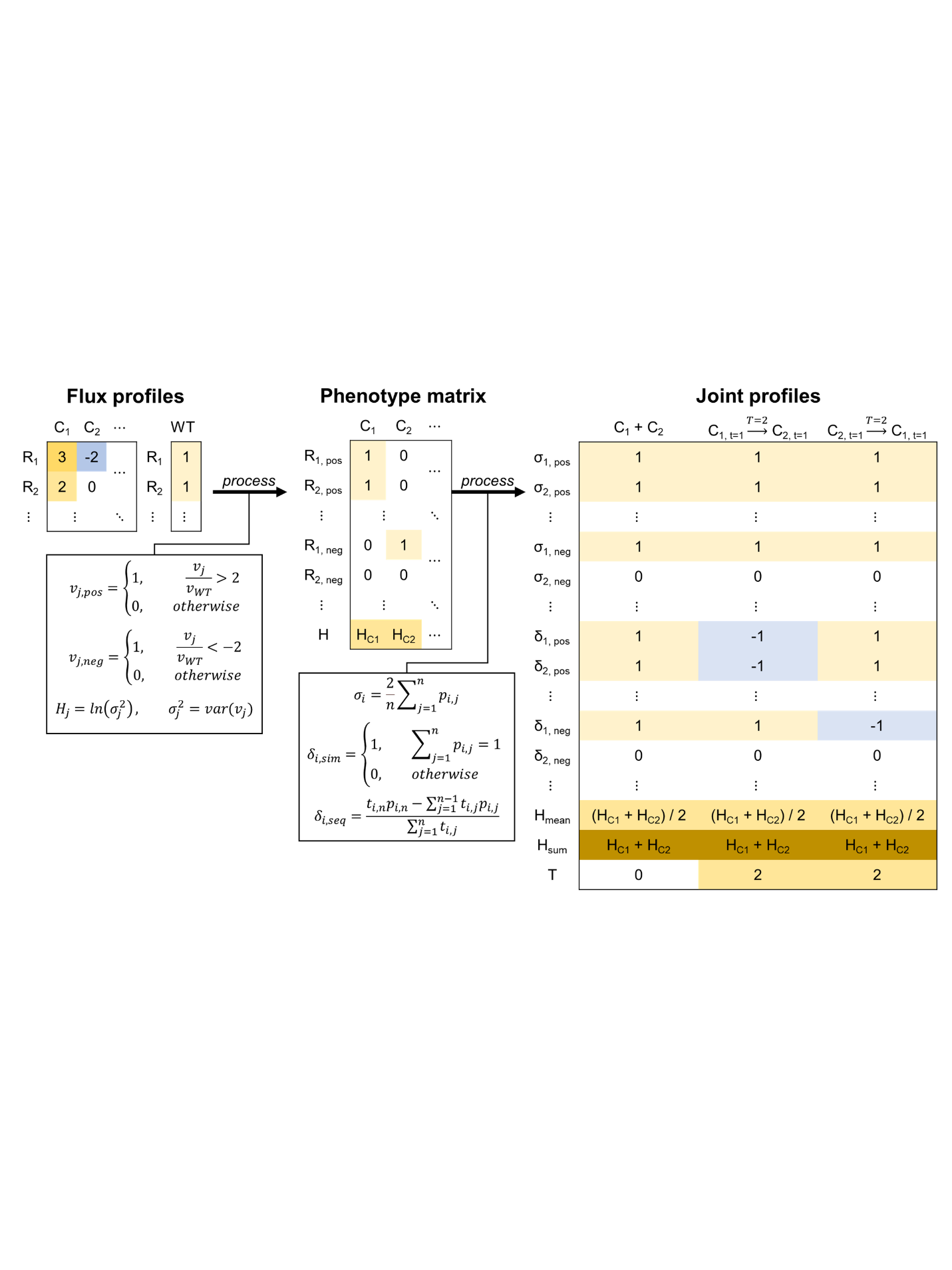

Fig. S2.

**Schematic of flux data processing into joint profiles.** Flux data (*v*) simulated from genome-scale metabolic models is binarized according to differential flux (either positive or negative) in comparison to wild type (WT, i.e., reference). These binarized flux profiles, along with the entropy (H) calculated for each condition (C), define the phenotype matrix which is subsequently processed into joint profiles. The sigma (σ) definition is the same between simultaneous (sim) and sequential (seq) interactions, while the delta (δ) definition differs depending on the interaction type. R = reaction, T = time interval, n = number of conditions in a combination.

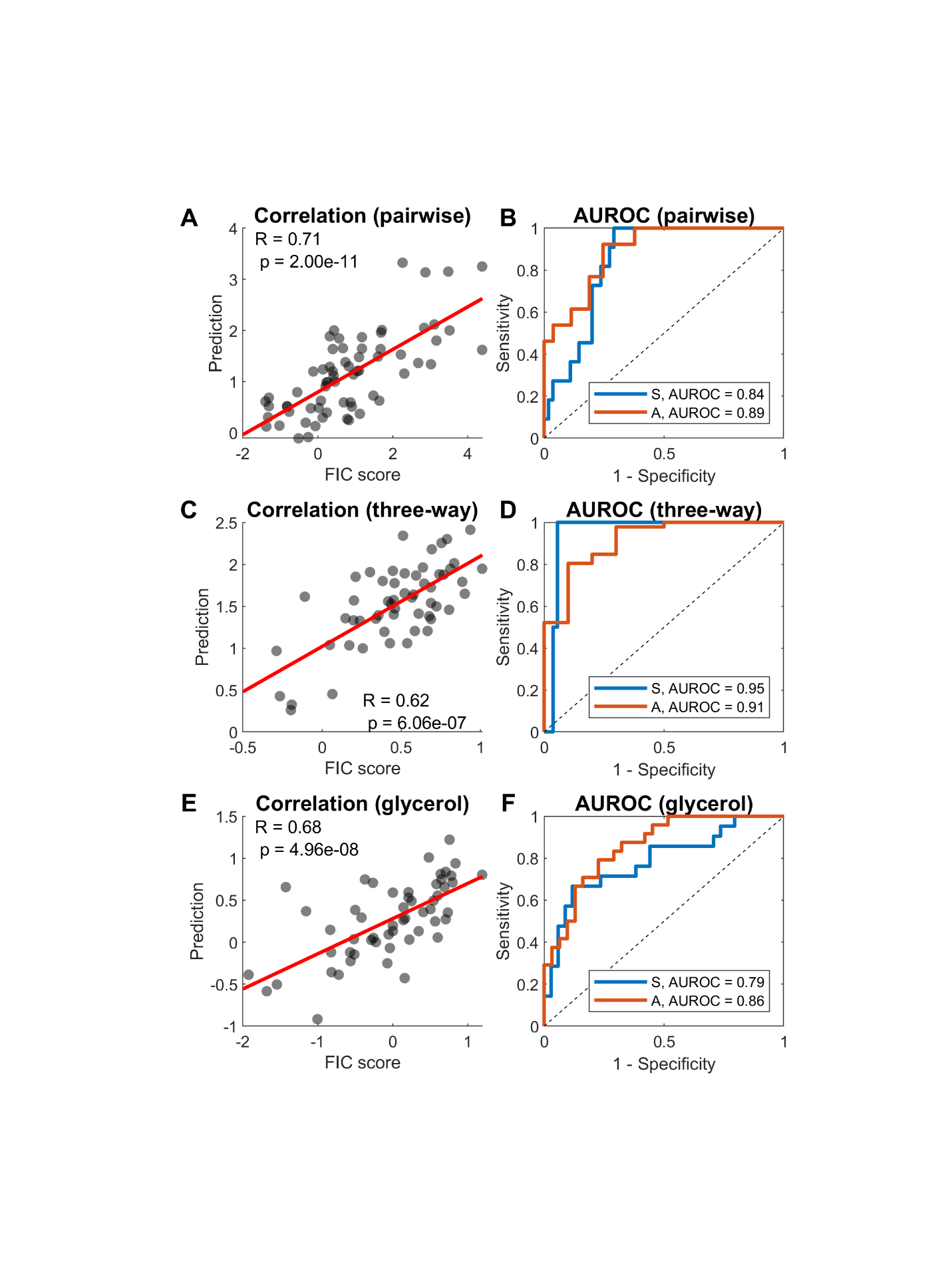

Fig. S3.

**CARAMeL results for *E. coli* drug interaction data.** Model performance results visualized as scatter and receiver operating curve (ROC) plots are shown for predicting (**A-B**) pairwise interactions, (**C-D**) three-way interactions, and (**E-F**) pairwise interactions in M9 glycerol. AUROC: area under the receiver operating curve. FIC: fractional inhibitory concentration, S: synergy, A: antagonism.

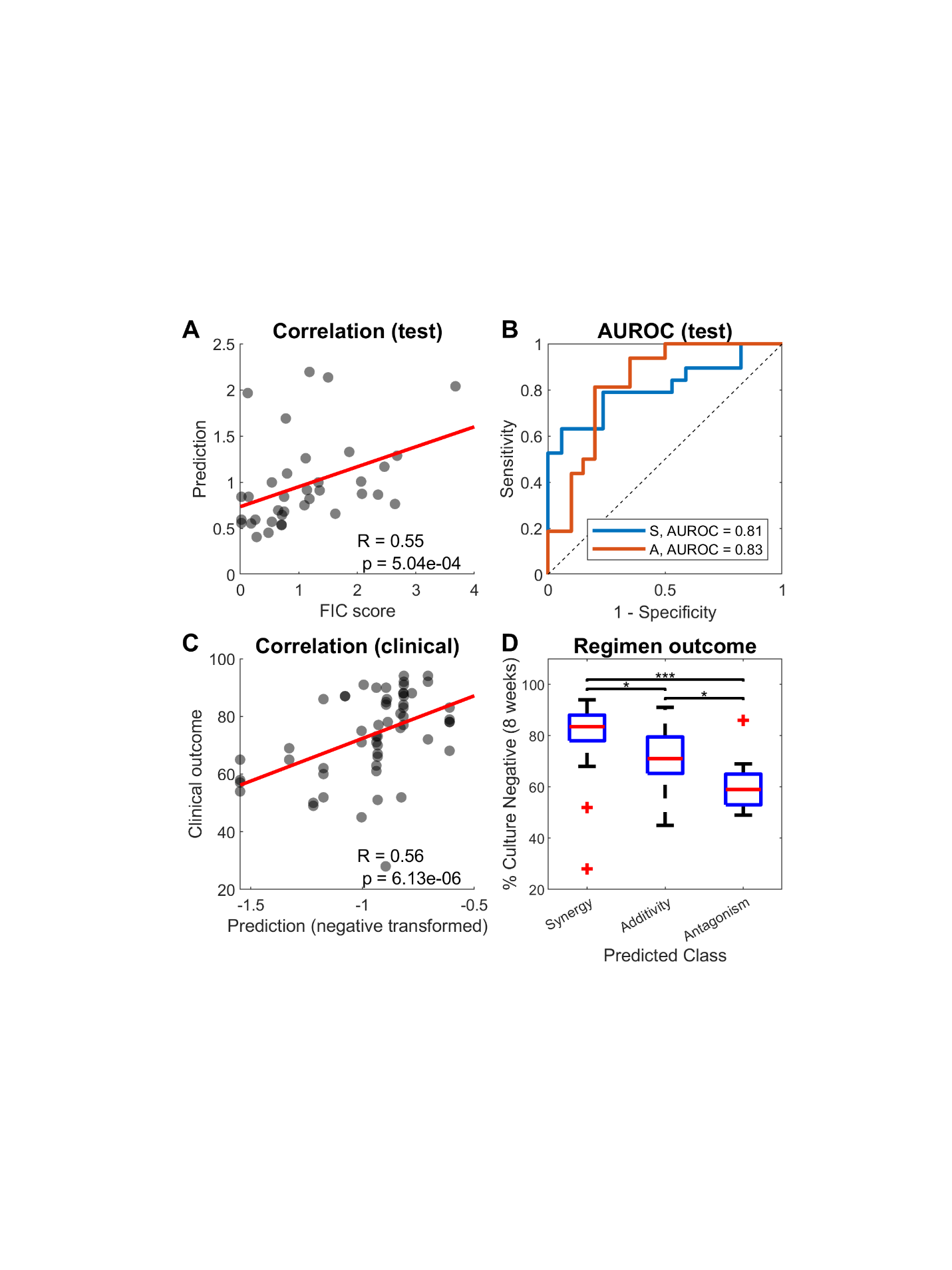

Fig. S4.

**CARAMeL results for M. tb drug interaction data.** (**A-B**) Model performance results visualized as scatter and receiver operating curve (ROC) plots are shown for predicting multi-drug interactions measured experimentally. (**C**) The model predictions for 57 TB regimens prescribed in clinical trials correlate with clinical efficacy. (**D**) Predictions classified as synergistic capture most of the efficacious treatments (sputum clearance > 80%). AUROC: area under the receiver operating curve, ** p-value < 0.01, *** p-value < 0.001 (unpaired t-test). FIC: fractional inhibitory concentration, AUROC: area under the receiver operating curve, S: synergy, A: antagonism.

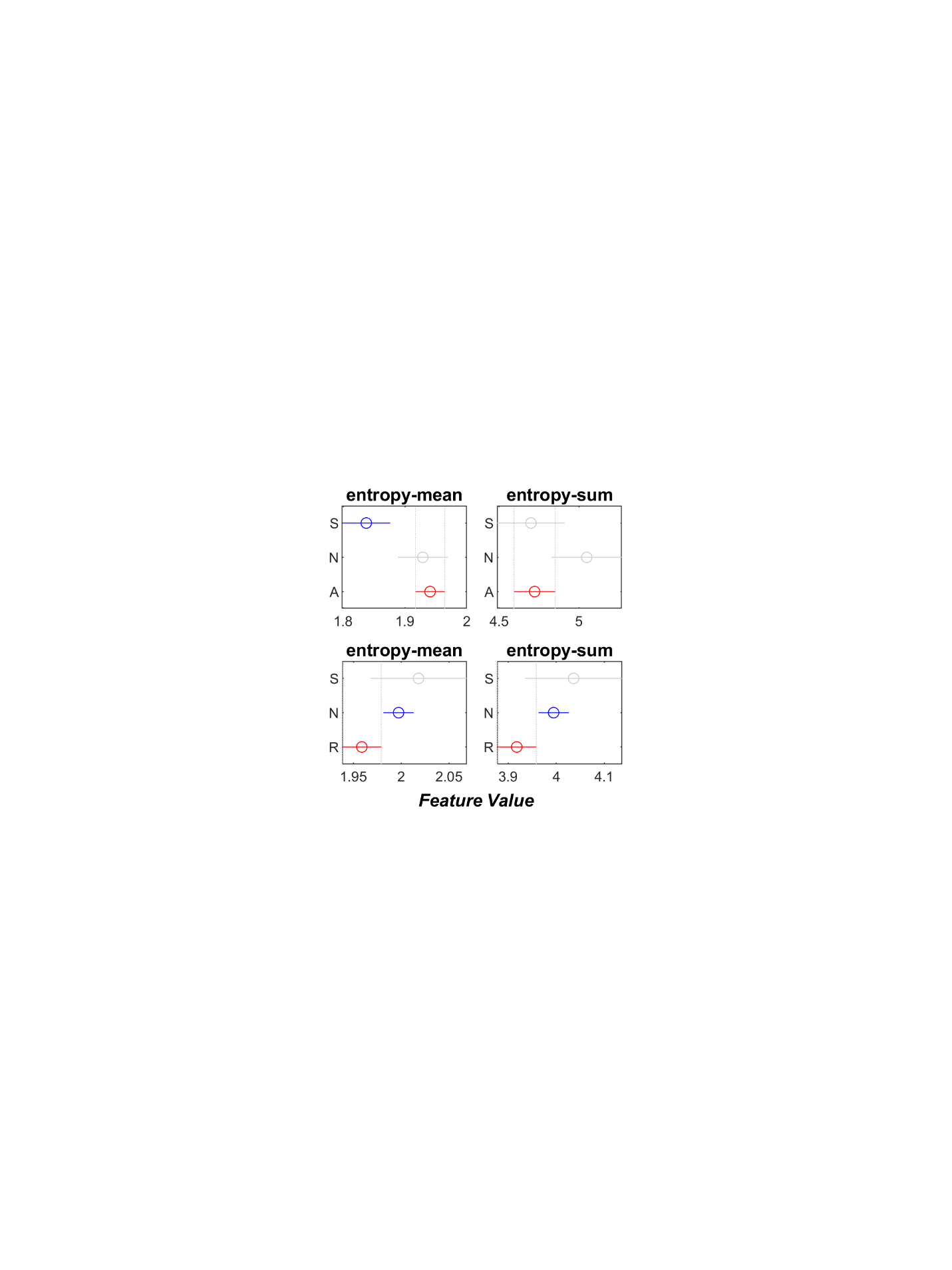

Fig. S5.

**Distribution of cell-specific drug interaction predictions.** The distribution for the top ten drug interactions with the largest variation across cells are shown for all time cases (D_1_ + D_2_, D_1_ → D_2_, D_2_ → D_1_). Refer to Tables S2 and S4 for full descriptions on antibiotics used for *E. coli*.

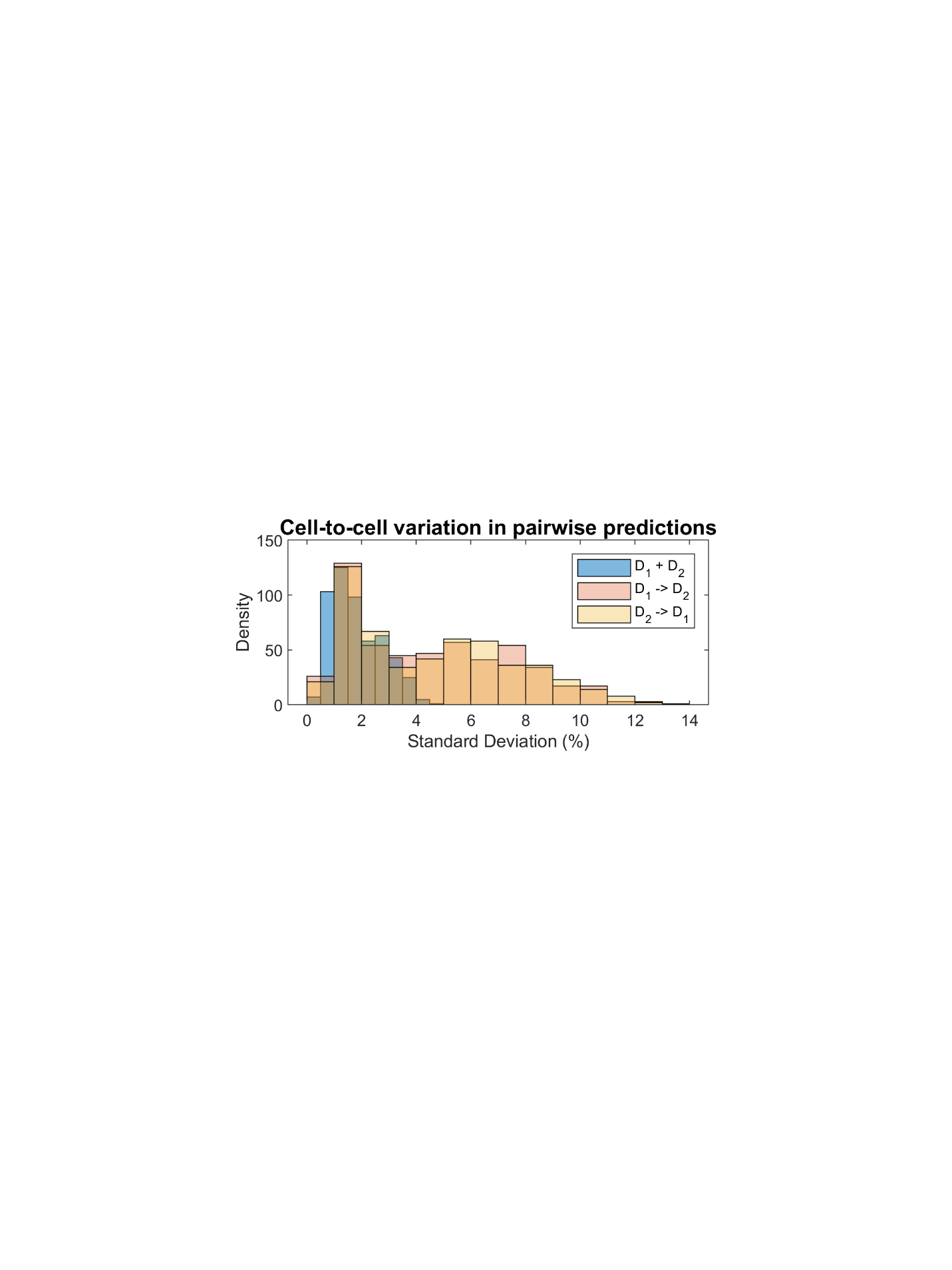

Fig. S6.

**Distribution of cell-specific drug interaction predictions.** The distribution for the top ten drug interactions with the largest variation across cells are shown for all time cases (D_1_ + D_2_, D_1_ → D_2_, D_2_ → D_1_). Refer to Tables S2 and S4 for full descriptions on antibiotics used for *E. coli*.

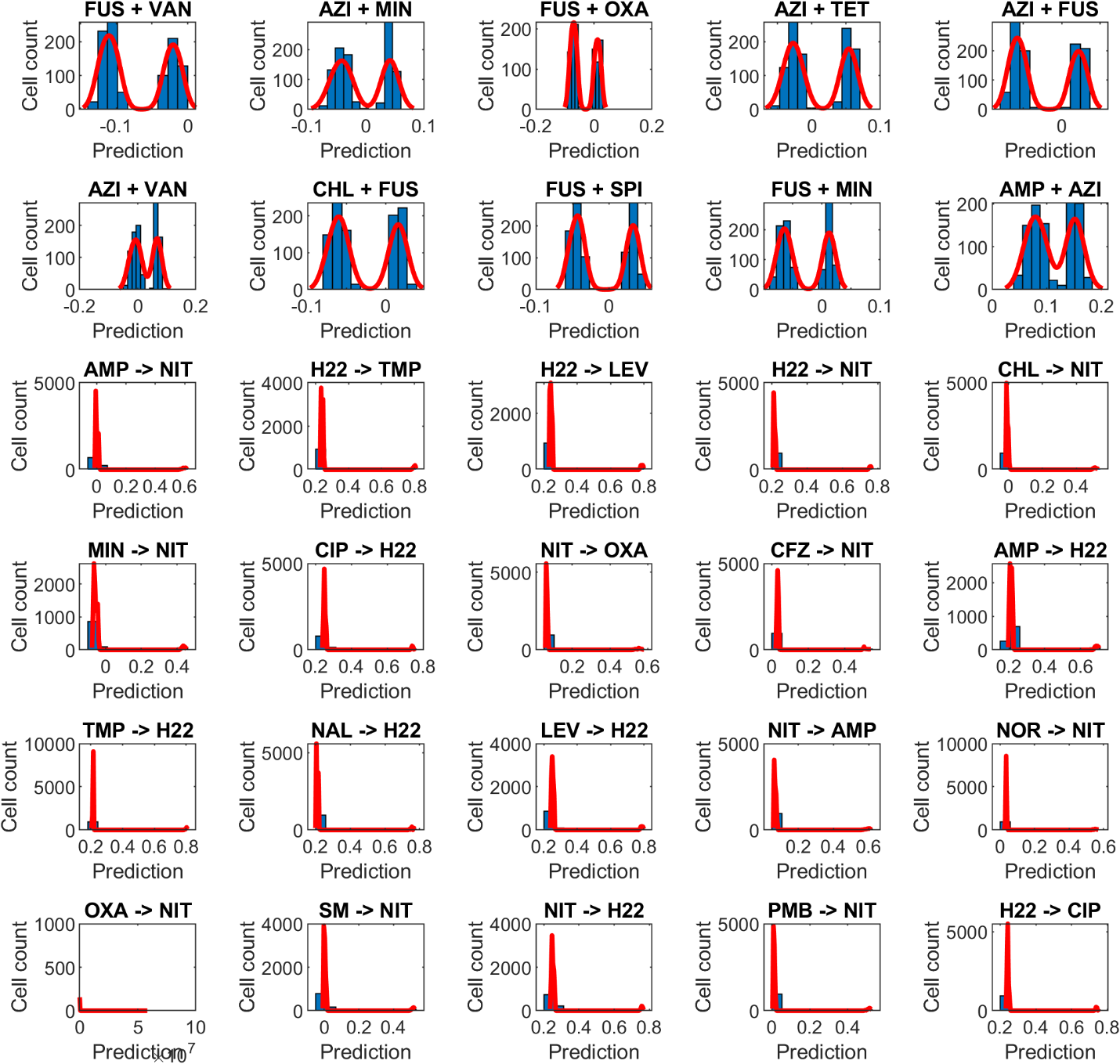

Fig. S7.

**Distribution of cell-specific drug interaction predictions.** The distribution for the top ten drug interactions with the largest variation across cells are shown for all time cases (D_1_ + D_2_, D_1_ → D_2_, D_2_ → D_1_). Refer to Tables S2 and S4 for full descriptions on antibiotics used for *E. coli*.

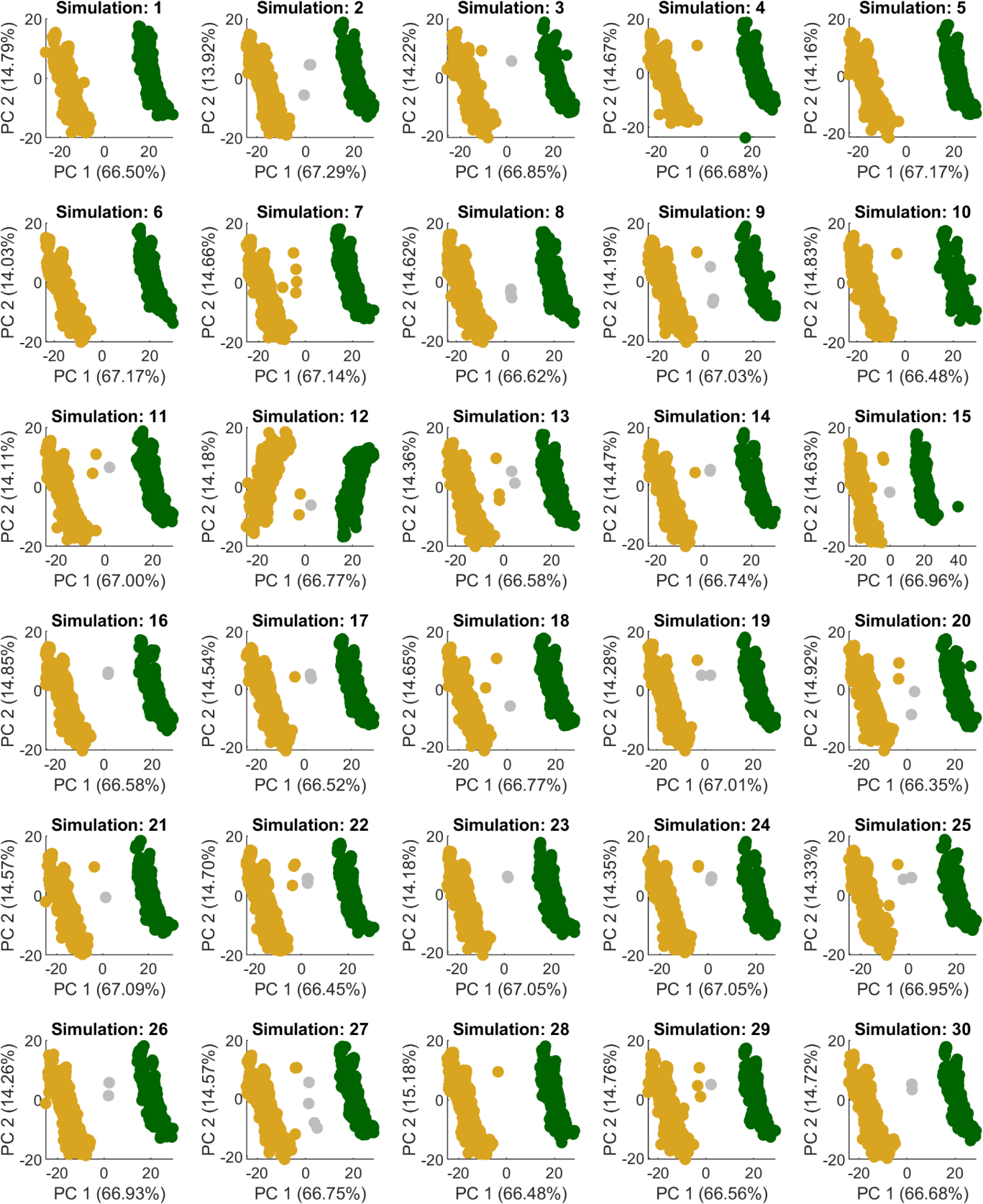

Fig. S8.

**Cell clustering based on drug interaction data is reproducible.** Repeated simulation (N = 30) of cell-specific fluxes (via population FBA) followed by CARAMeL prediction of cell-specific drug interaction outcomes reproducibly shows bimodal distribution of simultaneous prediction data, as seen in the 2-dimensional visualization of cell placement along the principal component (PC) space.

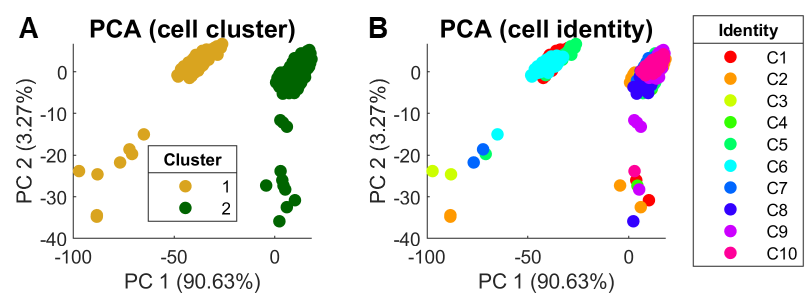

Fig. S9.

**Cell clustering is not driven by non-uniform sampling of the flux solution space.** Cell-specific drug interaction outcomes were repeatedly determined for a given cell based on uniform sampling of the flux solution space via optGpSampler. (**A**) The principal component analysis (PCA) visualization of the prediction data for simultaneous drug interactions shows similar clustering patterns as seen in Figs. 5B and S8. (**B**) The same PCA visualization grouped by cell identity shows that predictions belonging to the same original cell identity cluster together (except for a few outliers).

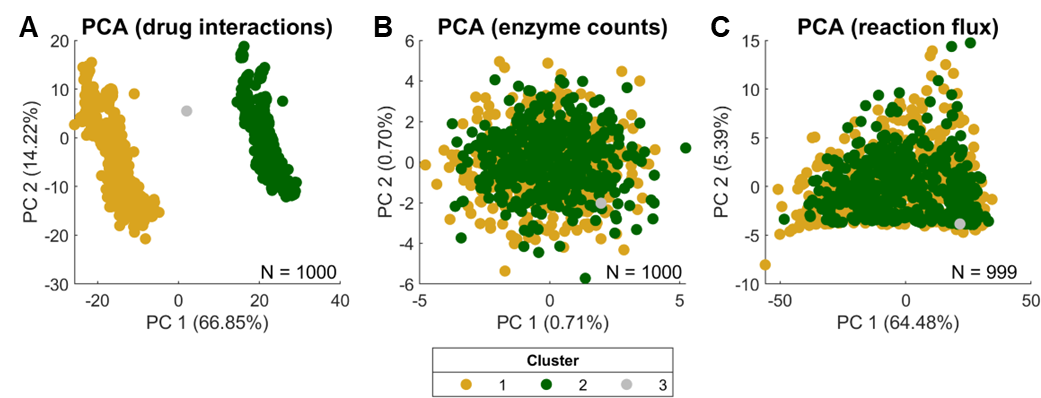

Fig. S10.

**Distinct cell clustering only occurs in CARAMeL prediction data for cell-specific drug interaction outcomes.** Cell clustering was investigated by applying principal component analysis (PCA) for cell-specific (**A**) drug interaction prediction data, (**B**) sampled enzyme count data, and (**C**) simulated metabolic flux data. The cluster groups shown in the legend were defined via k-means clustering of the PCA-transformed data for drug interactions. Of note, panel A is the same image shown in Fig. 5B in the main text. Additionally, panel C only shows the placement of 999 cells in PCA space due to removal of one outlier point.

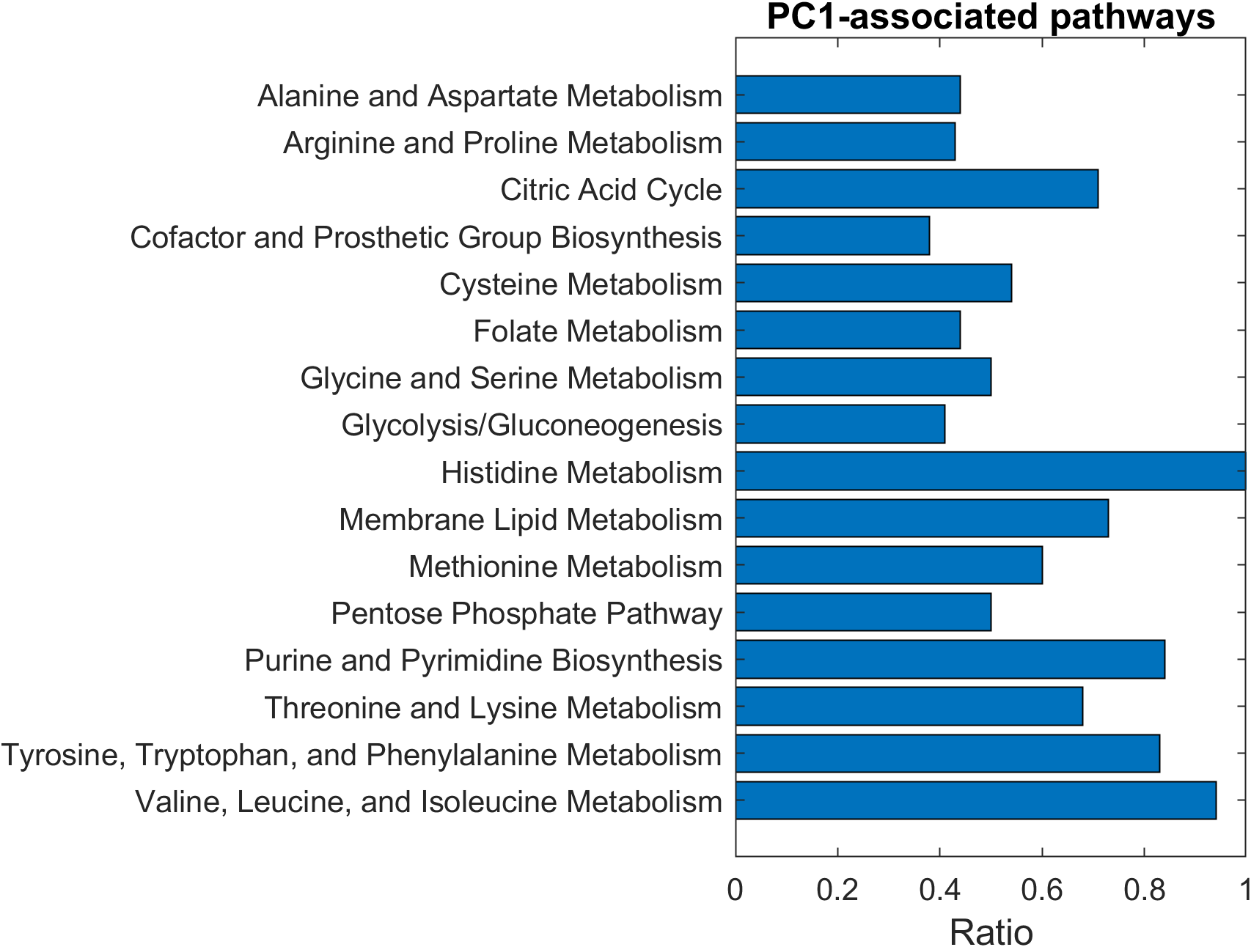

Fig. S11.

**16 metabolic pathways are significantly associated with cell clustering.** Over 400 metabolic reactions were found to robustly correlate in a significant manner with cell-specific scores along principal component 1 (PC1) shown in Fig. 5B (Data S7). A total of 16 pathways were subsequently found to be enriched by this set of reactions (hypergeometric test, adjusted p-value < 0.05).

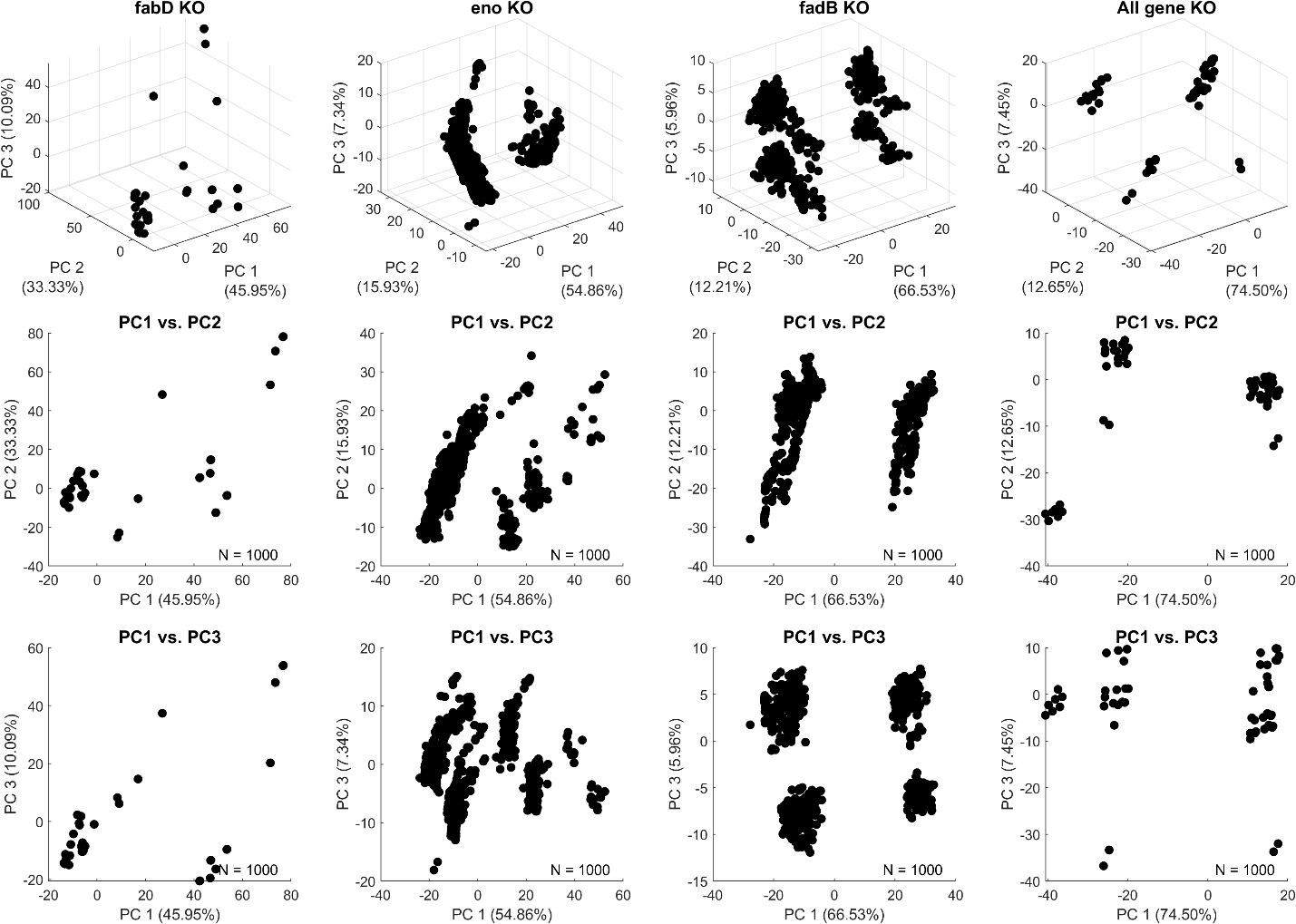

Fig. S12.

**Cell clustering is driven by stochastic changes in eno and fabD levels.** Single- and multi-gene knockout (KO) simulations for eno, fadB, and fabD reveal that the clustering pattern seen in Fig. 5B cannot be replicated in the absence of eno and fabD.

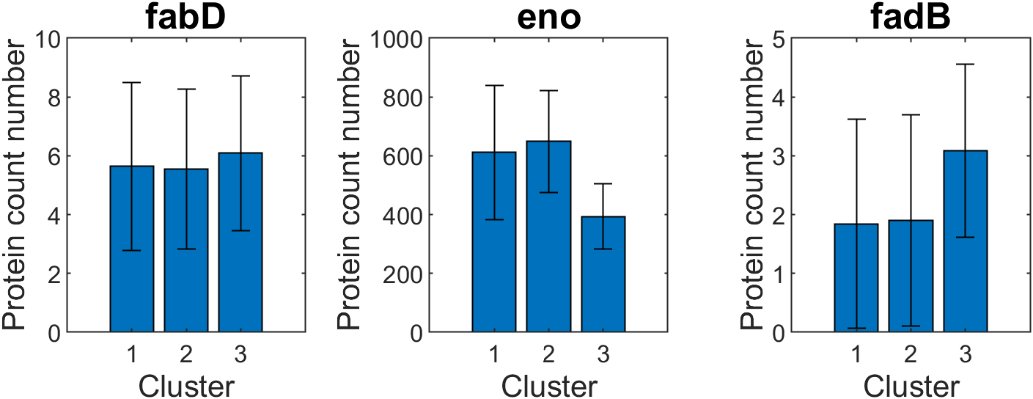

Fig. S13.

**Cluster-3 cells are characterized by low eno levels.** Cluster-3 cells, or those that do not cluster together with the dominant cell sub-populations seen in Fig. 5B, were found to possess much lower levels of eno (~400 per cell) compared to cluster-1 and cluster-2 cells (~600 per cell).

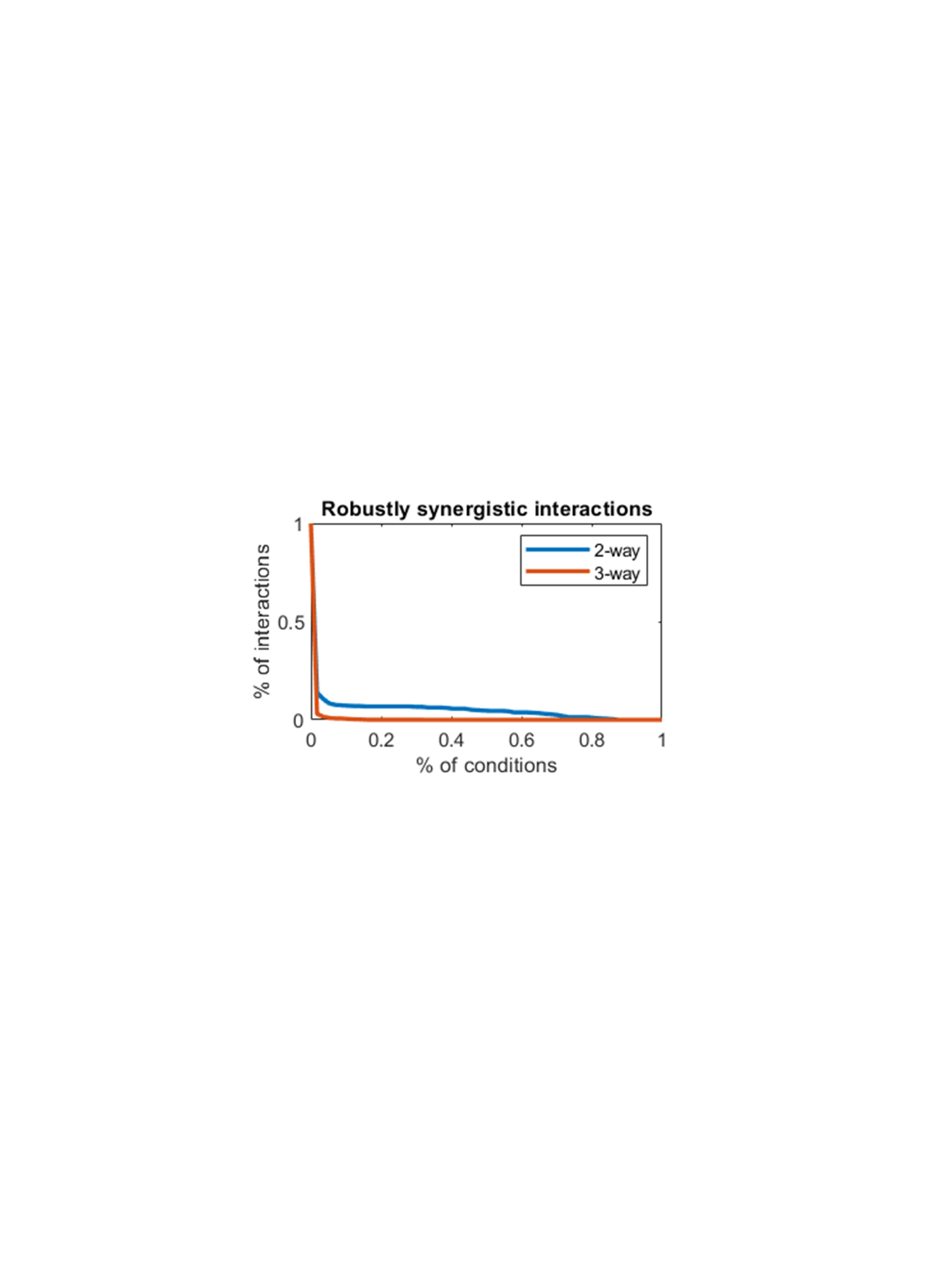

Fig. S14.

**Distribution of cell-specific drug interaction predictions.** The distribution for the top ten drug interactions with the largest variation across cells are shown for all time cases (D_1_ + D_2_, D_1_ → D_2_, D_2_ → D_1_). Refer to Tables S2 and S4 for full descriptions on antibiotics used for *E. coli*.

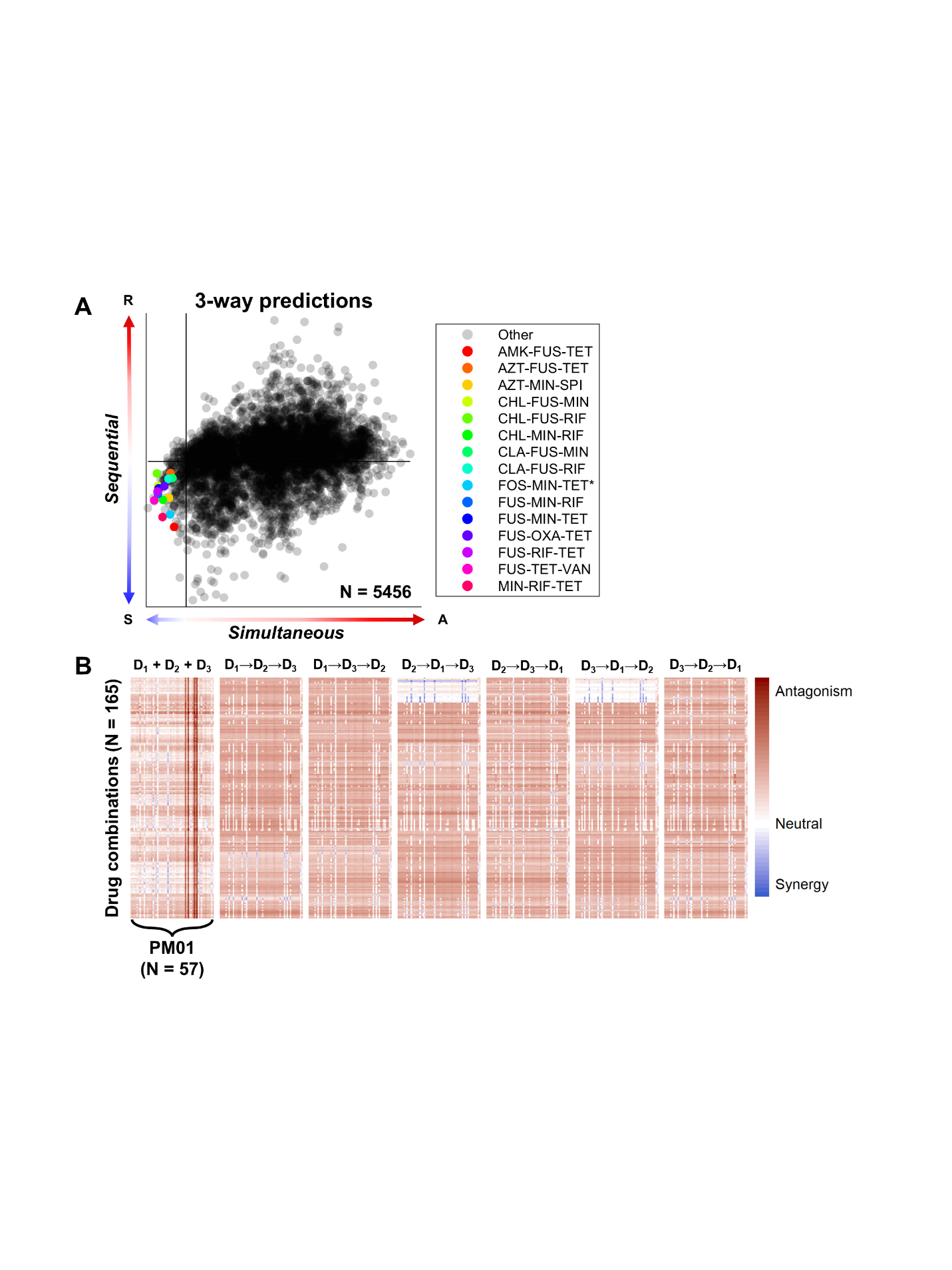

Fig. S15.

**CARAMeL predictions for three-way combination therapy landscape.** (**A**) Out of 5,456 unique three-way drug interactions, 165 were predicted to yield synergy (IS < 0) in at least one media condition for both the simultaneous and sequential cases (the top 15 robustly synergistic drug combinations are listed in the legend). (**B**) Heatmap of the predicted interaction scores for the 165 drug combinations across 57 media conditions and seven interaction types. Refer to Tables S2 and S4 for full descriptions on antibiotics used for *E. coli*.

Table S1.

**Omics-based approach evaluation for different z-score thresholds. *** Chosen parameters for *E. coli* and *M. tb*. R1: pairwise interactions (*36*), R2: three-way interactions (*37*), R3: glycerol interactions (*37*), R4: pairwise and three-way interactions (*38*, R5: pairwise to five-way TB clinical regimens (*55*).

|  | **Z-score** | ***E. coli* results** | | | ***M. tb results*** | |
| --- | --- | --- | --- | --- | --- | --- |
|  |  | *R1* | *R2* | *R3* | *R4* | *R5* |
|  | 0.25 | 0.1728 | 0.4548 | 0.5101 | 0.5604 | 0.4518 |
|  | 0.50 | 0.3158 | -0.0494 | 0.4004 | 0.5510 | 0.4870 |
|  | 0.75 | 0.2558 | 0.4535 | 0.6616 | 0.5283 | 0.3555 |
|  | 1.00 | 0.2780 | 0.3724 | 0.6781 | 0.5372 | 0.4802 |
|  | 1.25 | 0.3541 | 0.4384 | 0.6566 | 0.5637 | 0.4102 |
|  | 1.50 | 0.4269 | 0.5369 | 0.6501 | 0.5929 | 0.3889 |
|  | 1.75 | 0.3494 | 0.3891 | 0.6460 | 0.5234 | 0.4580 |
| ***** | 2.00 | 0.6508 | 0.5201 | 0.7068 | 0.5537 | 0.4737 |
|  | 2.25 | 0.4125 | 0.4671 | 0.6602 | 0.4489 | 0.4653 |
|  | 2.50 | 0.3665 | 0.4159 | 0.6432 | 0.4064 | 0.3937 |
|  | 2.75 | 0.3741 | 0.4621 | 0.6284 | 0.4139 | 0.4856 |
|  | 3.00 | 0.4194 | 0.3473 | 0.6177 | 0.4969 | 0.5344 |

Table S2.

**List of antibiotics used in *E. coli* drug interaction datasets.** Abb.: abbreviation.

|  |  |  |  | **Dataset** | | | | | |
| --- | --- | --- | --- | --- | --- | --- | --- | --- | --- |
| **Compound** | **Abb.** | **Target** | **Class** | **Pair (train)** | **Pair (test)** | **Three-way** | **LB** | **Glucose** | **Glycerol** |
| Amikacin | AMK | Protein synthesis, 30S | Aminoglycoside | ✓ | ✓ |  |  |  | ✓ |
| Gentamicin | GEN |  |  | ✓ | ✓ |  |  |  |  |
| Spectinomycin | SPE |  |  |  | ✓ |  | ✓ | ✓ | ✓ |
| Tobramycin | TOB |  |  | ✓ | ✓ |  |  |  |  |
| Minocycline | MIN |  | Tetracycline |  |  | ✓ |  |  |  |
| Tetracycline | TET |  |  | ✓ | ✓ |  | ✓ | ✓ | ✓ |
| Azithromycin | AZI | Protein synthesis, 50S | Macrolide |  |  | ✓ | ✓ | ✓ | ✓ |
| Chlarythromycin | CLA |  |  | ✓ | ✓ |  |  |  |  |
| Erythromycin | ERY |  |  | ✓ | ✓ |  |  |  |  |
| Chloramphenicol | CHL |  | Phenylpropanoid | ✓ | ✓ | ✓ | ✓ | ✓ | ✓ |
| Ciprofloxacin | CIP | DNA gyrase | Quinolone | ✓ | ✓ | ✓ |  |  |  |
| Levofloxacin | LEV |  |  | ✓ | ✓ |  |  |  |  |
| Nalidixic acid | NAL |  |  | ✓ | ✓ | ✓ |  |  | ✓ |
| Ampicillin | AMP | Cell wall | Beta-lactam |  |  | ✓ | ✓ | ✓ | ✓ |
| Aztreonam | AZT |  |  |  |  |  | ✓ | ✓ | ✓ |
| Cefoxitin | CEF |  |  | ✓ | ✓ |  |  |  | ✓ |
| Oxacillin | OXA |  |  | ✓ | ✓ |  |  |  |  |
| Vancomycin | VAN |  | Glycopeptide |  | ✓ |  |  |  |  |
| Fusidic acid | FUS | Elongation factor | Fusidane |  | ✓ |  |  |  |  |
| Trimethoprim | TMP | Folic acid biosynthesis | Pyrimidine | ✓ | ✓ |  |  |  |  |
| Rifampicin | RIF | RNA synthesis | Rifampin |  | ✓ | ✓ | ✓ | ✓ | ✓ |
| Nitrofurantoin | NIT | Multiple mechanisms | Furan | ✓ | ✓ | ✓ |  |  |  |
| Triclosan | TRI |  | Phenol |  |  |  | ✓ | ✓ | ✓ |
| Hydrogen peroxide | H22 | Oxidative stress | Stress | ✓ | ✓ |  |  |  |  |

Table S3.

**List of antibiotics used in *M. tb* drug interaction datasets.** Abb.: abbreviation, PTM: post-translational modification. * Putative mechanism.

|  |  |  |  | **Dataset** | | |
| --- | --- | --- | --- | --- | --- | --- |
| **Compound** | **Abb.** | **Target** | **Class** | **Train** | **Test** | **Clinical** |
| Sutezolid | SUTx | Protein synthesis, 23S | Oxazolidinone | ✓ |  |  |
| Amikacin | AMK | Protein synthesis, 30S | Aminoglycoside | ✓ |  |  |
| Kanamycin | KAN |  |  | ✓ |  |  |
| Spectinomycin | SPE |  |  | ✓ | ✓ |  |
| Streptomycin | SM |  |  | ✓ | ✓ | ✓ |
| Minocycline | MIN |  | Tetracycline | ✓ |  |  |
| Tetracycline | TET |  |  | ✓ |  |  |
| Azithromycin | AZI | Protein synthesis, 50S | Macrolide | ✓ |  |  |
| Chlarythromycin | CLA |  |  | ✓ | ✓ |  |
| Erythromycin | ERY |  |  | ✓ |  |  |
| Roxithromycin | ROX |  |  | ✓ |  |  |
| Linezolid | LZDx |  | Oxazolidinone | ✓ |  |  |
| Chloramphenicol | CHL |  | Phenylpropanoid | ✓ |  |  |
| Ciprofloxacin | CIP | DNA gyrase | Quinolone | ✓ | ✓ |  |
| Levofloxacin | LEV |  |  | ✓ | ✓ |  |
| Moxifloxacin | MOX |  |  | ✓ | ✓ | ✓ |
| Norfloxacin | NFX |  |  |  | ✓ |  |
| Ofloxacin | OFX1 |  |  | ✓ |  | ✓ |
| Novobiocin | NOV |  | Glycoside | ✓ |  |  |
| Ampicillin | AMP | Cell wall | Beta-lactam | ✓ |  |  |
| Oxacillin | OXA |  |  | ✓ |  |  |
| Vancomycin | VAN |  | Glycopeptide | ✓ |  |  |
| Cefaclor | CFL |  | Cephalosporin | ✓ |  |  |
| SQ109 | SQ109 |  | Ethylenediamine | ✓ |  |  |
| Isoniazid* | INH |  | Hydrazine | ✓ | ✓ | ✓ |
| Econazole | ECO |  | Imidazole | ✓ |  |  |
| Pretomanid | PA824 |  |  | ✓ |  | ✓ |
| Ethionamide* | ETH |  | Isonicotinic acid | ✓ |  |  |
| Cycloserine D | CSD |  | Serine | ✓ |  |  |
| PBTZ169 | PBTZ169x |  | Thiazine | ✓ |  |  |
| Capreomycin | CAP | Multiple mechanisms | Peptide | ✓ | ✓ |  |
| Clofazimine* | CFZ |  | Phenazine | ✓ | ✓ |  |
| Fusidic acid | FUS | Elongation factor | Fusidane | ✓ |  |  |
| Ethambutol | EMBx | RNA synthesis | Ethylenediamine | ✓ |  | ✓ |
| Rifampicin | RIF |  | Rifampin | ✓ | ✓ | ✓ |
| Rifapentine | RIFP |  | Rifamycin |  |  | ✓ |
| Bedaquiline | BDQ | ATP synthase | Diarylquinoline | ✓ | ✓ |  |
| Ethium bromide | EB | DNA structure | Phenanthridine | ✓ |  |  |
| Pyrazinamide* | PZA | Fatty acid synthase | Pyrazine |  |  | ✓ |
| Menadione | MEN | PTM | Vitamin | ✓ |  |  |
| Verapamil | VERx | Calcium channels | Phenethylamine | ✓ |  |  |
| Thioridazine | THZ | Synaptic activity | Phenothiazine | ✓ |  |  |
| Chlorpromazine | CPZ |  |  | ✓ | ✓ |  |

Table S4.

**List of antibiotics used in sequential drug interaction datasets for *E. coli*.** Abb.: abbreviation.

|  |  |  |  | **Time scale** | | |
| --- | --- | --- | --- | --- | --- | --- |
| **Compound** | **Abb.** | **Target** | **Class** | *T = 10* | *T = 21* | *T = 90* |
| Amikacin | AMK | Protein synthesis, 30S | Aminoglycoside | ✓ | ✓ | ✓ |
| Gentamicin | GEN |  |  | ✓ |  | ✓ |
| Spectinomycin | SPE |  |  |  | ✓ |  |
| Streptomycin | SM |  |  | ✓ | ✓ | ✓ |
| Tobramycin | TOB |  |  |  | ✓ |  |
| Doxycycline | DOX |  | Tetracycline |  | ✓ | ✓ |
| Minocycline | MIN |  |  | ✓ |  | ✓ |
| Tetracycline | TET |  |  | ✓ | ✓ | ✓ |
| Azithromycin | AZI | Protein synthesis, 50S | Macrolide | ✓ |  | ✓ |
| Erythromycin | ERY |  |  |  | ✓ |  |
| Spiramycin | SPI |  |  |  | ✓ |  |
| Chloramphenicol | CHL |  | Phenylpropanoid | ✓ | ✓ | ✓ |
| Ciprofloxacin | CIP | DNA gyrase | Quinolone | ✓ | ✓ | ✓ |
| Levofloxacin | LEV |  |  | ✓ |  | ✓ |
| Nalidixic acid | NAL |  |  | ✓ | ✓ | ✓ |
| Norfloxacin | NOR |  |  |  |  | ✓ |
| Ampicillin | AMP | Cell wall | Beta-lactam | ✓ | ✓ |  |
| Cefoxitin | CEF |  |  |  | ✓ |  |
| Ceftazidime | CFZ |  |  |  |  | ✓ |
| Amoxicillin | AMX |  |  | ✓ |  |  |
| Sulfamonomethoxine | SMM | Folic acid biosynthesis | Sulfonamide |  | ✓ |  |
| Trimethoprim | TMP |  | Pyrimidine | ✓ | ✓ | ✓ |
| Nitrofurantoin | NIT | Multiple mechanisms | Furan | ✓ | ✓ |  |
| Fosfomycin | FOS | Cell wall biogenesis | Phosphonic acid | ✓ |  |  |
| Fusidic acid | FUS | Elongation factor | Fusidane |  | ✓ |  |
| Polymyxin B | PMB | Lipopolysaccharide | Peptide | ✓ |  |  |
| Rifampicin | RIF | RNA synthesis | Rifampin | ✓ |  | ✓ |

Table S5.

**Drug information for Biolog experiment.** Abb.: abbreviation, Conc.: drug concentration.

|  |  |  |  |  | **Conc. (μg/mL)** | |
| --- | --- | --- | --- | --- | --- | --- |
| **Compound** | **Abb.** | **Target** | **Class** | **Type** | *Single* | *Pairwise* |
| Aztreonam | AZT | Cell wall | Beta-lactam | Bactericidal | 0.03 | - |
| Cefoxitin | CEF |  |  | Bactericidal | 1.87 | 1.87 |
| Tetracycline | TET | Protein synthesis, 30S | Tetracycline | Bacteriostatic | 1.42 | 1.42 |
| Tobramycin | TOB |  | Aminoglycoside | Bactericidal | 0.15 | 0.15 |

Table S6.

**Constraint-based modeling (CBM) parameter optimization results.** ***** Chosen parameters for *M. tb*, **+** chosen parameters for *E. coli*. CV-R: 10-fold cross-validation correlation in the training dataset, GR-V: variance in growth rate, NG-P: percentage of no growth (GR = 0) conditions.

|  | **CBM parameters** | | | ***E. coli* results** | | | ***M. tb* results** | | |
| --- | --- | --- | --- | --- | --- | --- | --- | --- | --- |
|  | *Kappa* | *Rho* | *Epsilon* | *CV-R* | *GR-V* | *NG-P* | *CV-R* | *GR-V* | *NG-P* |
|  | 0.001 | 0.001 | 0.001 | 0.4055 | 0 | 0 | 0.3640 | 0 | 0 |
|  | 0.01 | 0.01 | 0.001 | 0.3260 | 0.0074 | 0 | 0.5024 | 0.0003 | 0.0233 |
|  | 0.1 | 0.1 | 0.001 | 0.4077 | 0.0634 | 0 | 0.4858 | 0.0005 | 0.0698 |
|  | 1 | 1 | 0.001 | 0.4313 | 0.2022 | 0.3636 | 0.4409 | 0.0005 | 0.0698 |
|  | 0.001 | 0.001 | 0.01 | 0.4260 | 0 | 0 | 0.4124 | 0 | 0 |
| ***** | **0.01** | **0.01** | **0.01** | 0.3739 | 0.0091 | 0 | 0.5207 | 0.0003 | 0.1395 |
|  | 0.1 | 0.1 | 0.01 | 0.389 | 0.0670 | 0 | 0.5149 | 0.0004 | 0.6977 |
|  | 1 | 1 | 0.01 | 0.4189 | 0.1866 | 0.4848 | 0.5242 | 0.0003 | 0.7674 |
|  | 0.001 | 0.001 | 0.1 | 0.6406 | 0 | 0 | 0.5231 | 0 | 0 |
|  | 0.01 | 0.01 | 0.1 | 0.4090 | 0.0095 | 0 | 0.4959 | 0.0001 | 0.5581 |
|  | 0.1 | 0.1 | 0.1 | 0.3978 | 0.1056 | 0.0606 | 0.4792 | 0.0001 | 0.7907 |
|  | 1 | 1 | 0.1 | 0.3869 | 0.1294 | 0.5152 | 0.4471 | 0.0001 | 0.8837 |
|  | 0.001 | 0.001 | 1 | 0.3860 | 0 | 0 | 0.4620 | 0 | 0 |
| **+** | **0.01** | **0.01** | **1** | 0.6512 | 0.0113 | 0 | 0.5131 | 0.0003 | 0.6047 |
|  | 0.1 | 0.1 | 1 | 0.6150 | 0.0994 | 0.0909 | 0.5270 | 0 | 0.8140 |
|  | 1 | 1 | 1 | 0.6294 | 0.0765 | 0.5758 | 0.5099 | 0 | 0.9070 |

Table S7.

**Benchmarking correlation results based on different constraint-based modeling (CBM) parameter choices.** ***** Chosen parameters for *M. tb*, **+** chosen parameters for *E. coli*. R1: pairwise interactions (*36*), R2: three-way interactions (*37*), R3: glycerol interactions (*37*), R4: pairwise and three-way interactions (*38*), R5: pairwise to five-way TB clinical regimens (*55*).

|  | **CBM parameters** | | | ***E. coli* results** | | | ***M. tb results*** | |
| --- | --- | --- | --- | --- | --- | --- | --- | --- |
|  | *Kappa* | *Rho* | *Epsilon* | *R1* | *R2* | *R3* | *R4* | *R5* |
|  | 0.001 | 0.001 | 0.001 | 0.2884 | 0.4441 | 0.5781 | 0.6370 | 0.5535 |
|  | 0.01 | 0.01 | 0.001 | 0.5032 | 0.3516 | 0.5092 | 0.5873 | 0.4369 |
|  | 0.1 | 0.1 | 0.001 | 0.4676 | 0.3666 | 0.5715 | 0.4946 | 0.2304 |
|  | 1 | 1 | 0.001 | 0.5115 | 0.4001 | 0.5421 | 0.4717 | 0.4135 |
|  | 0.001 | 0.001 | 0.01 | 0.3726 | 0.3636 | 0.5544 | 0.4638 | 0.5361 |
| ***** | **0.01** | **0.01** | **0.01** | 0.4525 | 0.2447 | 0.5900 | 0.5256 | 0.5445 |
|  | 0.1 | 0.1 | 0.01 | 0.5390 | 0.2599 | 0.5514 | 0.4858 | 0.2642 |
|  | 1 | 1 | 0.01 | 0.3899 | 0.3460 | 0.5915 | 0.4730 | 0.4124 |
|  | 0.001 | 0.001 | 0.1 | 0.5421 | 0.5809 | 0.5669 | 0.5382 | 0.4483 |
|  | 0.01 | 0.01 | 0.1 | 0.5829 | 0.5023 | 0.6281 | 0.6335 | 0.4263 |
|  | 0.1 | 0.1 | 0.1 | 0.3313 | 0.3762 | 0.6382 | 0.5942 | 0.5140 |
|  | 1 | 1 | 0.1 | 0.1545 | 0.4375 | 0.6745 | 0.5560 | 0.4982 |
|  | 0.001 | 0.001 | 1 | 0.1772 | 0.1372 | 0.2668 | 0.3253 | 0.5380 |
| **+** | **0.01** | **0.01** | **1** | 0.6445 | 0.6216 | 0.6641 | 0.4884 | 0.4870 |
|  | 0.1 | 0.1 | 1 | 0.6057 | 0.6536 | 0.6650 | 0.4947 | 0.4788 |
|  | 1 | 1 | 1 | 0.6352 | 0.6169 | 0.6101 | 0.4939 | 0.3726 |

Data S1 to S10 (Available at <https://www.dropbox.com/s/859ebsx1drri1xc/data.xlsx?dl=0>)

Data S1. (separate file)

Top CARAMeL features explaining 95% of the variance in model predictions.

Data S2. (separate file)

iJO1366 reactions explaining 95% of the variance between actual and predicted interaction outcomes.

Data S3. (separate file)

Biolog phenotype microarray 1 (PM01) conditions.

Data S4. (separate file)

CARAMeL predictions for 2-way drug interactions for 1,000 individual cells and three treatment strategies (1 x simultaneous, 2 x sequential) (N = 1,584,000).

Data S5. (separate file)

Cell-to-cell variation in predictions for pairwise drug interaction outcomes (N = 528).

Data S6. (separate file)

Principal component (PC) loadings 1 and 2 for the principal component analysis (PCA) transformation of the drug interaction prediction data (simultaneous only).

Data S7. (separate file)

iJO1366 reactions that are robustly* associated with cell clustering. *Significant correlation with PC1 scores determined for all 30 replicated runs.

Data S8. (separate file)

Cluster-based sensitivity and tolerance indication for the 528 unique drug pairs.

Data S9. (separate file)

CARAMeL predictions for 2-way drug interactions in 57 media conditions and three treatment strategies (1 x simultaneous, 2 x sequential) (N = 90,288).

Data S10. (separate file)

CARAMeL predictions for 2-way drug interactions in 57 media conditions and seven treatment strategies (1 x simultaneous, 6 x sequential) (N = 2,176,944).
